# Insulin synthesis is sustained by Tent5 poly(A) polymerases

**DOI:** 10.64898/2026.07.04.736497

**Authors:** Ana Kotte, Fabio Marcuccio, Linda Masante, Nadja von Wiegen, Lucia Prandi, Raquel Sofia Silva, Rodrigo Pracana, Jacopo Zasso, Blagoje Soskic, Alessandra Zappulo, Ivano Legnini

**Affiliations:** Human Technopole, Viale Rita Levi-Montalcini 1, 20157, Milan, Italy

**Author notes:** Equal contribution (Ana Kotte, Fabio Marcuccio).

**Keywords:** Insulin, mRNA metabolism, beta cells, TENT5, poly(A) tail, Type 2 Diabetes

## Abstract

Insulin is an essential regulator of glucose homeostasis in vertebrates, and impairment of its synthesis or action leads to diabetes with severe health complications in humans. It is therefore essential to understand how beta cells control insulin synthesis and secretion, including the transcription, translation and decay of its messenger RNA.

Using sequencing-based poly(A) tail length profiling from human tissue, genetic evidence for type 2 diabetes, bulk and single-cell transcriptomics and perturbation experiments, here we find that the insulin mRNA is stabilized by the activity of noncanonical poly(A) polymerases of the Tent5 family. We show that Tent5 activity is specific, promoted by both localization at the endoplasmic reticulum and regulatory sequences within the insulin mRNA and regulated by glucose.

Overall, our findings provide a mechanistic link between the dynamic control of insulin production by beta cells and the direct regulation of insulin mRNA metabolism.

## Introduction

The insulin mRNA is extremely abundant, with estimates up to 150,000 molecules per cell accounting for almost a third of the total mRNA pool in pancreatic beta cells (1). Given its central role in glucose metabolism, insulin synthesis and release are highly dynamic and must be tightly regulated at both the mRNA and protein levels. In fact, glucose-dependent control of insulin mRNA transcription, translation and degradation have been extensively reported (2–27), with a yet incomplete understanding of the underlying regulatory logic. For these reasons, dissecting the regulation of insulin mRNA metabolism is essential for understanding beta cell function and glucose control, including their impairment in diabetes.

Mammalian mRNAs are synthesized in the nucleus and decorated with a poly(A) tail of approximately 250 nucleotides (28,29), which is required for translation initiation (30). Once exported into the cytoplasm, mRNAs are deadenylated at a gene-specific rate (31–33). This rate is a major determinant of mRNA stability, since deadenylation precedes and is required for canonical mRNA degradation (34,35). Recently, it has been shown that poly(A) tails of specific mRNAs can be elongated in the cytoplasm by noncanonical poly(A) polymerases such as the TErminal Nucleotidyl Transferase 5 family proteins (TENT5A-D in humans, previously known as FAM46A-D) (36–38). Interestingly, TENT5 activity has been linked to secretory function by specific targeting of endoplasmic reticulum (ER)-residing mRNAs, including for example Immunoglobulin mRNAs in B cells and vaccine mRNAs in macrophages (39,40).

Type 2 Diabetes (T2D) is characterized by a failure of beta cells to compensate for insulin resistance, leading to chronic hyperglycaemia with severe health outcomes (41). Beta cells respond to elevated glucose in prediabetes by increasing proinsulin synthesis, which in turn causes and exacerbates ER stress (15,22). In fact, misfolding of proinsulin at the ER of beta cells and the consequent triggering of the Unfolded Protein Response (UPR) is one of the first molecular hallmarks of diabetes (42). An emerging yet substantial body of evidence therefore suggests ER stress as a key pathogenetic mechanism of T2D, and targeting UPR regulators as well as insulin synthesis itself has shown beneficial results in animal models of diabetes, thus representing a potential novel strategy to intercept T2D (43–48). Interestingly, TENT5C expression was found to cause ER stress-mediated cell death in multiple myeloma (49–51). Altogether, these observations suggest that TENT5 family proteins may participate in cellular responses to ER stress through the regulation of mRNA metabolism of secretory proteins.

Given the high secretory demand of pancreatic beta cells and the tight coupling between insulin biosynthesis and ER homeostasis, we hypothesized that TENT5 proteins may regulate insulin mRNA metabolism under physiological conditions and potentially play a role in the pathogenesis of T2D. In this work, we leverage third-generation sequencing-based poly(A) tail length profiling, bulk and single-cell transcriptomics data from human pancreatic islets, *in vitro* beta cell models and perturbation experiments to study the role of TENT5 proteins in insulin mRNA regulation in physiological and pathological conditions.

## Results

### 1. Poly(A) tail length profiling reveals that mRNAs encoding for pancreatic endocrine hormones carry unusually long tails

To gather a high-throughput, genome-wide characterization of poly(A) tail length distributions in the human pancreas, we generated and processed Full-length, Poly(A) tail and mRNA sequencing (FLAM-seq(52)) from three independent samples. The resulting combined dataset contains 4,492,513 single-molecule, real-time sequencing reads, including 1,691,379 high-quality reads, defined by the presence of unique cDNA library adapters and a bona fide poly(A) tail (Fig. 1A). Of these, 636,361 span full-length transcript sequences, extending from the 5’ CAP to the poly(A) tail (Fig. 1B and Suppl. Fig. 1A-B). These reads were assigned to 34,870 unique transcripts from 19,215 genes (Fig. 1A and Suppl. Fig 1B).

**1.**
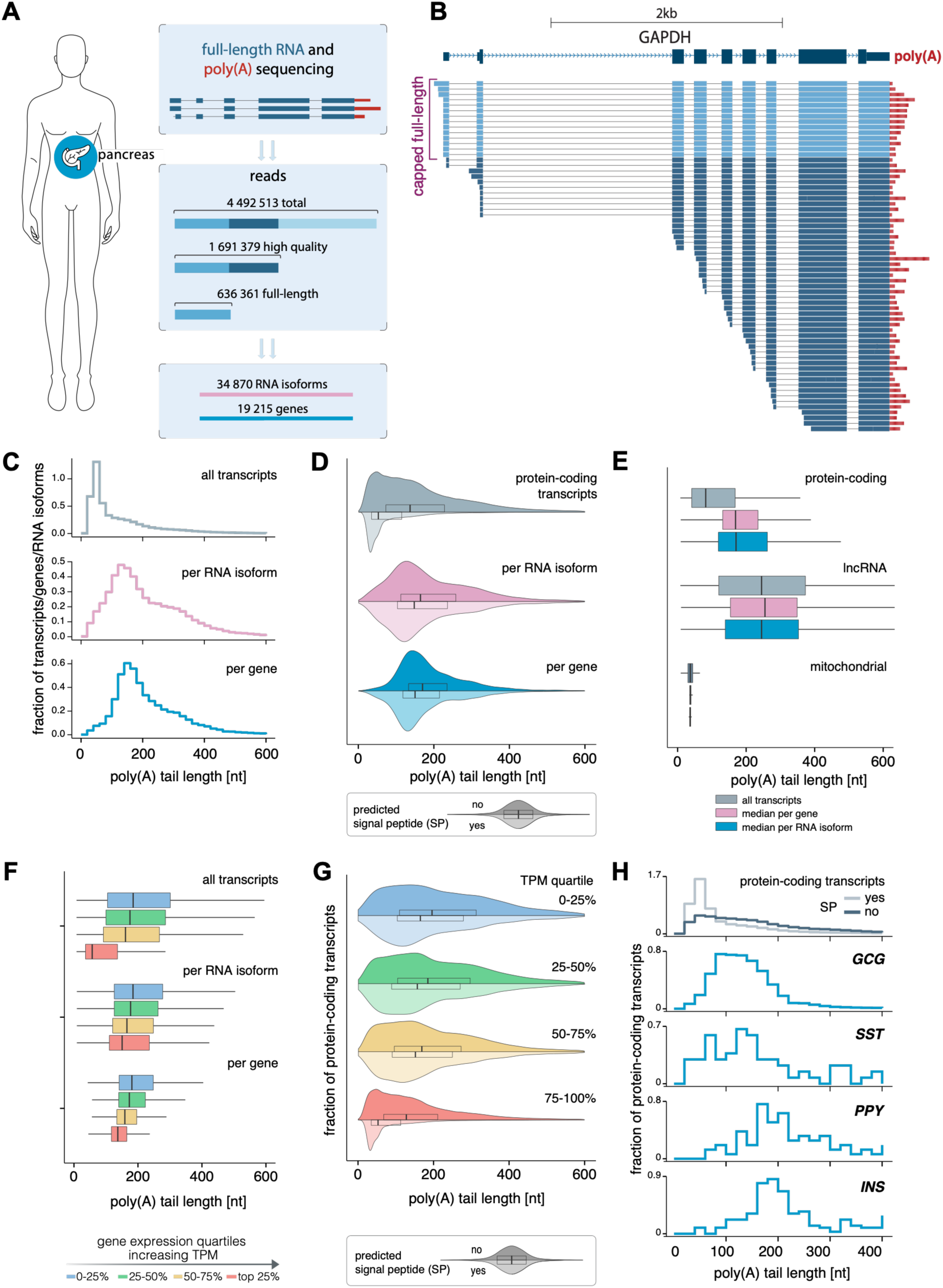
A subset of secretory mRNAs has unusually long poly(A) tails. A. Schematic overview of sequencing statistics for the combined human pancreas FLAMseq dataset (n=3 independent RNA samples and datasets). Numbers indicate the total number of sequencing reads, the number of high-quality reads with unique cDNA adaptors and a bona fide poly(A) tail and the number of full-length reads spanning entire RNA isoforms, including the 5’cap. B. Browser shot example of read distributions and their poly(A) tails for *GAPDH* in a FLAMseq pancreas dataset. Highlighted are full-length reads spanning the entire gene, including the 5’ cap. For simplicity only a subset of forward aligned reads is displayed. C. Poly(A) tail length distributions in the combined human pancreas dataset (n=3 independent RNA samples and datasets) across all detected transcripts (top) or summarized as median poly(A) tail length per RNA isoform (middle) or gene (bottom). D. Poly(A) tail length distributions of protein-coding genes in the combined human pancreas dataset, stratified by the absence (upper violin) or the presence (lower violin) of a predicted or validated signal peptide in at least one RNA isoform of the corresponding gene, according to SignalP6.0(110). Distributions across all detected transcripts or summarized as median poly(A) tail length per gene and per RNA isoform are shown. E. Boxplots of poly(A) tail lengths in nuclear encoded protein-coding genes, long non-coding RNA (lncRNA) and mitochondrial genes across all detected transcripts or summarized as median poly(A) tail length per gene and per RNA isoform. F. Boxplots of poly(A) tail lengths for genes stratified into equal expression quartiles based on transcripts per million (TPM) values obtained from the combined FLAMseq dataset. Poly(A) tail lengths are shown either across all detected transcripts or as median values summarized at the gene or RNA isoform level. G. Poly(A) tail length distributions for protein-coding transcripts in the combined human pancreas dataset, binned into equal expression quartiles based on transcript per million (TPM) values obtain from FLAMseq and stratified by the absence (upper violin) or presence (lower violin) of a predicted or validated signal peptide in at least one RNA isoform of the corresponding gene, according to SignalP 6.0(110). H. Poly(A) tail length distributions of the indicated pancreatic hormones compared to all other protein-coding genes within the same gene expression quartile (upper expression bin 75-100% quartile). Protein coding genes are stratified by the absence or the presence of a predicted or validated signal peptide (SP) in at least one RNA isoform of the corresponding gene, according to SignalP 6.0(110)

We observed high tail length variability among genes and transcripts, with a peak at 34 nt, a median at 69 nt and an interquartile range comprised between 38 nt and 161 nt (Fig. 1C and Suppl. Fig. 1C), similarly to what has been previously observed in human cells (52), and suggesting gene and transcript-specific RNA metabolism control. Tail lengths per gene were reproducible across replicates, with increasing correlation as a function of RNA abundance, as previously observed (52) (Suppl. Fig. 1D).

Given the recent observation of mRNAs encoding for secreted proteins being re-adenylated by TENT5 enzymes (53), we stratified protein-coding genes by the presence of a validated or predicted signal peptide. We did not observe generally longer poly(A) tails for secretory mRNAs but the opposite (Fig. 1D and Suppl. Fig. 1E). Instead, gene and transcript-level variability were dominated by other factors such as transcript class, with mitochondrial RNAs carrying short poly(A) tails and long non-coding RNAs having considerably longer tails (Fig. 1E and Suppl. Fig. 1F), similarly to what has been previously observed in human cell lines (52,54). Moreover, we observed decreasing tail length distributions for increasing RNA abundance quartiles, also in accordance with previous observations (Fig. 1F and Suppl. Fig. 1G) (52,54). This trend was consistent for both mRNAs carrying a signal peptide and all other mRNAs (Fig. 1G and Suppl. Fig. 1H).

Despite signal peptide-encoding mRNAs and abundant mRNAs carrying shorter tails on average, we observed that mRNAs encoding for the major pancreatic endocrine hormones such as insulin (*INS*), glucagon (*GCG*), somatostatin (*SST*) and pancreatic polypeptide (*PPY*) possess significantly longer tails than the bulk of protein-coding transcripts (Fig. 1H), albeit *SST* and *PPY* were detected with fewer reads in two of the replicates (Suppl. Fig. 1I). For the insulin mRNA in particular, we measured a mean poly(A) tail length across replicates larger than 99.5% of all mRNAs with comparable abundance in the human pancreas.

In summary, to the best of our knowledge, we provide the first long-read based compendium of transcript isoforms in the human pancreas together with isoform-level poly(A) tail length measurements, which revealed an unusually long poly(A) tail distribution for a subset of specific secretory mRNAs encoding for endocrine hormones, and particularly for the insulin mRNA.

### 2. TENT5A and C are highly expressed in pancreatic beta cells and TENT5C is genetically associated with T2D

Given the recently reported involvement of TENT5 family enzymes in stabilizing secretory mRNAs, we examined the expression of their encoding genes across human tissues in FANTOM (55,56) and GTEx (57) data. We observed that the *TENT5A* and *C* mRNAs are particularly abundant in the human pancreas, where we found unusually long poly(A) tails for endocrine hormone-encoding mRNAs (Fig. 2A).

**2.**
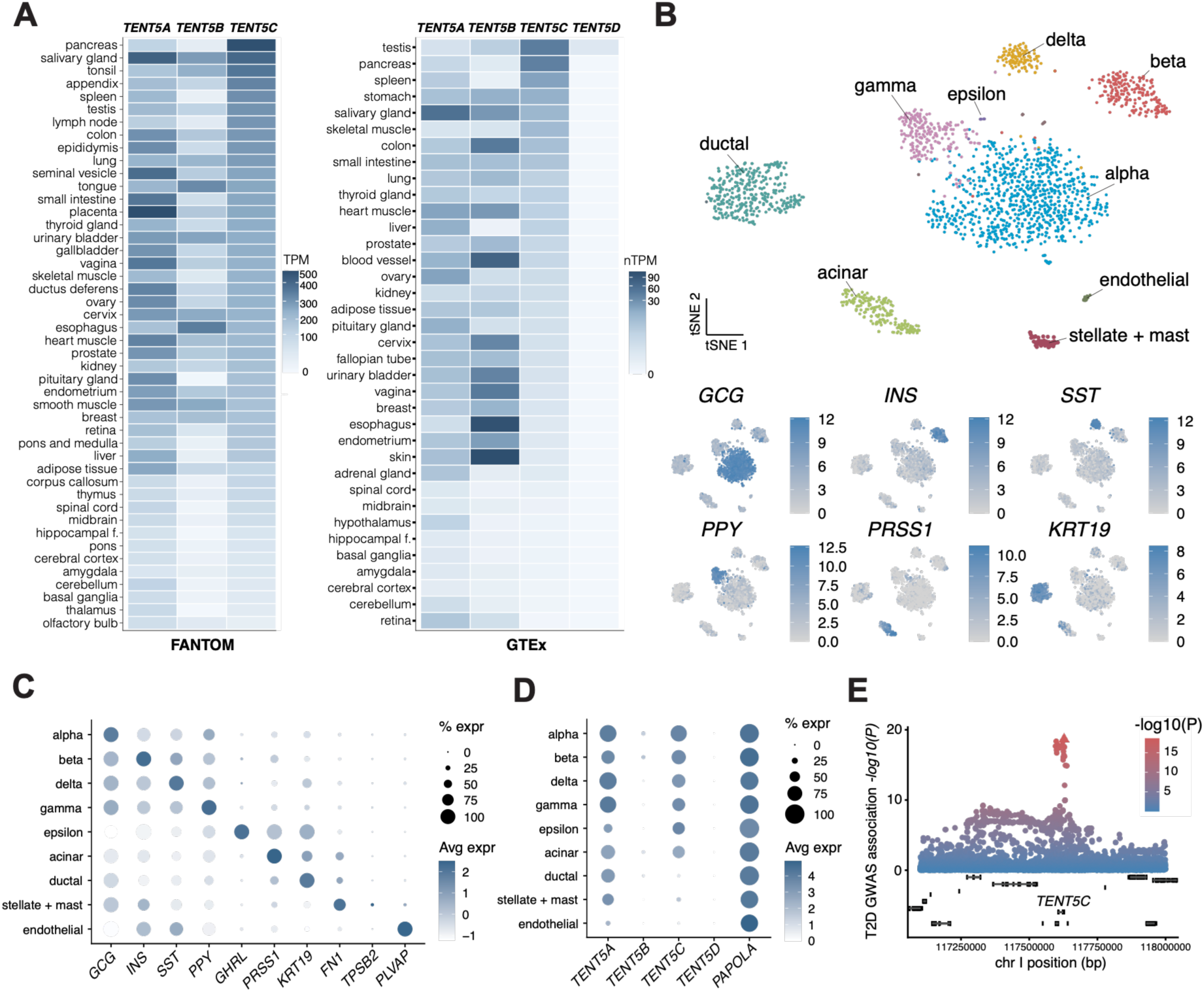
TENT5C is highly expressed in pancreatic beta cells and genetically associated with T2D. A. *TENT5 family* expression across human tissues as reported in FANTOM5 (55,56) and GTEx (57). FANTOM5 data are shown as scaled tags per million (TPM) per tissue, while GTEx data are presented as normalized protein coding transcripts per million (nTPM) across donors. *TENT5D* was not detected in any FANTOM5 datasets. Data available from human protein atlas (v25.1.proteinatlas.org). B. t-SNE projection for the dataset E-MTAB-5061 single-cell transcriptomes coloured by annotated cell type identity. Two additional datasets shown in Suppl. Fig. 2A. Distinct clusters correspond to endocrine populations (Alpha, Beta, Gamma, Delta, Epsilon), exocrine cells (Acinar and Ductal), endothelial, and mesenchymal/immune (Stellate + Mast) populations. C. Dot plot showing the scaled average expression of canonical marker genes across pancreatic cell types. Endocrine populations (Alpha, Beta, Gamma, Delta, Epsilon) are defined by *GCG*, *INS*, *SST*, *PPY*, and *GHRL* expression, whereas *PRSS1* and *KRT19* mark acinar and ductal cells, respectively, and *FN1/TPSB2* and *PLVAP* define mesenchymal/immune and endothelial clusters. D. Dot plot showing the average expression of *TENT5* family genes (*TENT5A-D*) and PAPOLA across pancreatic cell types. E. GWAS association with T2D at the *TENT5C* locus on chromosome I. Triangle indicates the highest value of association.

To investigate the cell type specificity of *TENT5* gene expression, we re-analyzed three single-cell RNA-seq datasets obtained from human pancreas (58,59). After sequencing, read processing, dimensionality reduction and clustering, we could retrieve all the expected pancreatic cell types (Fig. 2B and Suppl. Fig. 2A-B). These included endocrine alpha, beta, gamma, delta, epsilon cells, and exocrine acinar and ductal cells, as indicated by the expression of well-established cell-type specific markers and particularly by the expression of the genes encoding for the hormones secreted by the pancreatic endocrine cell types (Fig. 2C and Suppl. Fig. 2C). We next examined *TENT5* gene expression in all cell types and found that *TENT5A/C* are both expressed in islets, with *TENT5A* being more abundant across cell types and *TENT5C* more specifically enriched in endocrine cells, particularly in alpha and beta cells, whereas other *TENT5* paralogues were only poorly expressed (Fig. 2D and Suppl. Fig. 2D). Notably, *TENT5C* was more abundant than *TENT5A* in matched human islets bulk RNA-seq data (58), which generally provide more robust abundance estimates across genes than single-cell RNA-seq (60,61) (Suppl. Fig. 2E), as well as in a larger bulk RNA-seq dataset (62,63) (Suppl. Fig. 2E). Conversely, based on both bulk and single-cell data, *TENT5B* and *D* are barely expressed or absent in pancreatic islets (Fig. 2D and Suppl. Fig. 2E).

Based on these observations, we next examined the genetic association between *TENT5A/C*, T2D, and glycaemic traits. Interrogation of the Open Targets database (64) identified a T2D-associated locus with lead variant *1_117627844_CTG_C* (65), for which *TENT5C* was assigned a high L2G score of causality (L2G = 0.912). Notably, the lead fine-mapped variant is located within the 3′ Untranslated Region (UTR) of *TENT5C* (Fig. 2E). We observed consistent associations also in GWAS of blood glucose levels (Suppl. Fig. 2F), while we found no strong associations between *TENT5A* and traits associated with T2D.

We therefore re-analyzed two islets bulk RNA-seq datasets, one matched with the single cell-RNA-seq data in Fig. 2B-E and obtained from 3 healthy individuals and 4 T2D donors (58), and a larger one, from 33 healthy and 10 T2D donor islets (62,63). In both datasets we observed a slight reduction in *TENT5C* mRNA abundance, consistent with the loss of beta cells in T2D patient islets (66), and as indicated by a similar reduction in the insulin mRNA abundance (Suppl. Fig. 2G). Of note, when instead looking at *TENT5C* expression within single beta cells, not only was it not reduced, but instead slightly, albeit not significantly, increased in T2D compared to healthy donors in all three datasets, similarly to *TENT5A* (Suppl. Fig. H).

In summary, *TENT5A-C* are highly expressed in pancreatic beta cells and *TENT5C* is genetically associated with glycaemia and T2D. Our results so far led us to hypothesize that TENT5A-C might regulate beta cell function and insulin mRNA metabolism.

### 3. Insulin mRNA abundance is controlled by Tent5a/c

To test our hypothesis, we transfected the rat insulinoma cell line INS-1E with a pool of siRNAs targeting *Tent5a/c*^1^. After 48 hours, we collected RNA and measured *Tent5a/*c abundance via qRT-PCR, observing a 64% depletion of *Tent5c* upon RNAi and a 34% depletion in *Tent5a* (Fig. 3A). Following *Tent5a/c* depletion, we observed a reduction in the insulin mRNAs *Ins1* and *Ins2* of 54% and 58% respectively (Fig. 3A), resulting in reduced insulin secretion (Suppl. Fig. 3A). Knock-down of *Tent5a* alone was also sufficient to reduce *Ins1* and *Ins2* mRNA abundance, while that of *Tent5c alone* was sufficient to reduce *Ins2* mRNA abundance (Suppl. Fig. 3B-3C).

**3.**
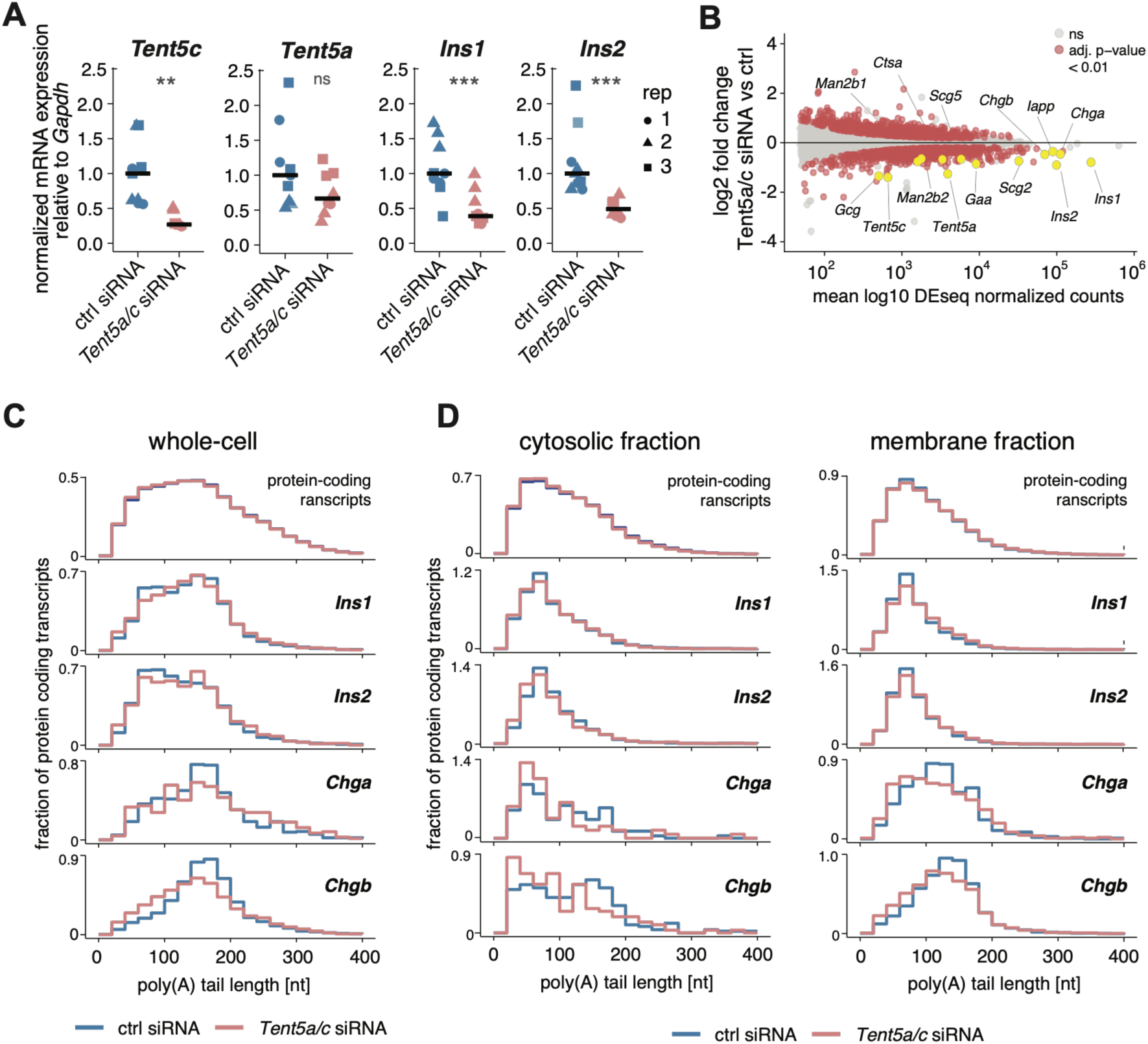
Insulin mRNA abundance is controlled by Tent5a/c poly(A) polymerases. A. qRT-PCR analysis of the indicated genes in INS-1E control cells (ctrl siRNA) and *Tent5a/c* double-knockdown cells *(Tent5a/c* siRNA). Expression was normalized to *Gapdh* and to control cells. Data represent n = 3 independent experiments, each with technical triplicates; crossbars indicate medians. Paired t-test p-values indicated as follows: <0.01 (**), <0.001 (***), ns (non-significant). B. Differential gene expression in INS-1E cells following double *Tent5a/c* knockdown (n=3, independent experiments). MA plot displaying differential expression as log2 fold change (*Tent5a/c* siRNA vs ctrl siRNA) versus mean DESeq2-normalized expression. Genes with adjusted p-value < 0.01 are highlighted in red, while non-significant genes are shown in grey. Selected significantly differentially expressed genes of interest are highlighted in yellow. Differential expression analysis was performed using DESeq2; genes with mean normalized counts < 50 were excluded prior to testing. C. Poly(A) tail length distributions of all protein coding transcripts or for the indicated genes in INS-1E control cells (ctrl siRNA) or *Tent5a/c* double knockdown cells (*Tent5a/c* siRNA). Distributions are normalized so that the area under each curve equals 1. Aggregated results across n=3 independent experiments are shown. D. Poly(A) tail length distributions of all protein coding transcripts or for the indicated genes in cytosolic or membrane subcellular fractions for INS-1E control cells (ctrl siRNA) or Tent5a/c double knockdown cells (*Tent5a/c* siRNA). Distributions are normalized so that the area under each curve equals 1. Aggregated results across n=3 independent experiments are shown.

In order to characterize the global effects of *Tent5a/c* on gene expression, we performed bulk RNA-seq on three independent replicates of control and *Tent5a/c* siRNA-treated INS-1E cells. Differential gene expression analysis confirmed strong and significant downregulation of *Tent5a*, *Tent5c,* as well as *Ins1 and Ins2.* In addition to those, we observed significant downregulation of mRNAs encoding for additional hormone peptides or factors involved in hormone peptide packaging such as *Gcg* (Glucagon), *Iapp* (Islet amyloid polypeptide), *Scg2/3/5* (Secretogranins), *Chga*, *Chgb* (Chromogranins), and a variety of lysosomal enzymes (e.g. *Man2b1/2*, *Ctsa*, *Gaa*) (Fig. 3B). To measure the effect of *Tent5a/c* depletion on poly(A) tail length, we performed FLAM-seq on bulk RNA samples from three independent replicates of control and *Tent5a/c* siRNA-treated INS-1E cells, where we did not observe a substantial shortening of insulin mRNA poly(A) tail length (Fig. 3C and Suppl. Fig. 3E). We hypothesized that a direct effect of *Tent5a/c* depletion on steady state tail length distributions might be masked by the coexistence of distinct subcellular pools of the insulin mRNAs and therefore performed FLAM-seq on purified membrane-enriched and cytosolic mRNA fractions from control and *Tent5a/c-*depleted INS1E cells (fraction purity in Suppl. Fig. 3D). We did not observe *Ins1/2* tail shortening in the membrane fraction, while we observed a mild shortening in the cytosolic one, especially for *Ins2* (Fig. 3D and Suppl. Fig. 3F,G). Of note, Silva, Mayrhofer et al. recently reported similar observations in total RNA and proposed that tail shortening might result in rapid mRNA degradation, thus masking the effect of Tent5 depletion on tail length at steady state (67). Conversely, for some of the most abundant and strongly downregulated genes upon *Tent5a/c* knock-down such as *Chga* and *Chgb*, we did detect tail shortening in both membrane and cytosolic fractions at steady state (Fig. 3D).

In summary, we found that TENT5 proteins control insulin mRNA abundance and secretion, as well as abundance and poly(A) tail length of mRNAs encoding for factors involved in insulin packaging such as chromogranins.

### 4. Tent5-mediated insulin mRNA regulation is stimulated by both its 3’ UTR and ER localization

TENT5 proteins were shown to localize at the ER surface by docking onto transmembrane proteins of the FNDC3 family (49), and to specifically re-adenylate mRNAs encoding for secreted proteins (40,53). Conversely, we observed that mRNAs encoding for secreted proteins do not have longer poly(A) tails than other mRNAs in general, even in human pancreatic tissue (Fig. 1D and Suppl. Fig. 1E), leading us to hypothesize that additional specificity factors underlying TENT5 proteins activity may be in place. Before testing this hypothesis, we sought to confirm Tent5 proteins subcellular localization in INS-1E cells. After trying a number of commercially available antibodies which resulted in unspecific signal (s. Methods and Suppl. Fig. 4A), we purified INS1-E subcellular fractions (Fig. 4A, Suppl. Fig.4B) and performed label-free quantification (LFQ)-based proteomics (Fig. 4B). Only peptides uniquely mapping to *Tent5a* were detected among *Tent5* family members. *Tent5a* showed a modest but significant enrichment in the membrane fraction relative to the cytoplasmic fraction (log2FC = 0.874, adjusted p-value = 0.0025, Fig. 4B) and to the total lysate (log2FC = 0.588, adjusted p-value = 0.0021), which is compatible with this protein being enriched at the ER but also suggesting a substantial *Tent5a* fraction existing in the cytosol. To then test whether *Tent5a/c* activity is specific to mRNAs targeted to the secretory pathway, we stratified the *Tent5a/c* knock-down bulk RNA-seq data (Fig. 3B) by the presence of a signal peptide in protein coding genes. Despite only a fraction of mRNAs encoding signal peptide-containing proteins being downregulated, genes in this category were substantially enriched among differentially expressed genes (p-value = 2.8e-30) and were almost exclusively downregulated (Fig. 4C). In fact, gene set enrichment analysis revealed “Protein processing in the ER” and related gene sets as the top enriched pathways amongst downregulated genes (Fig. S4 C-D). However, we did not observe significant poly(A) tail shortening in genes encoding for signal peptide containing proteins following *Tent5a/c* knock-down (Suppl. Fig. 4E).

**4.**
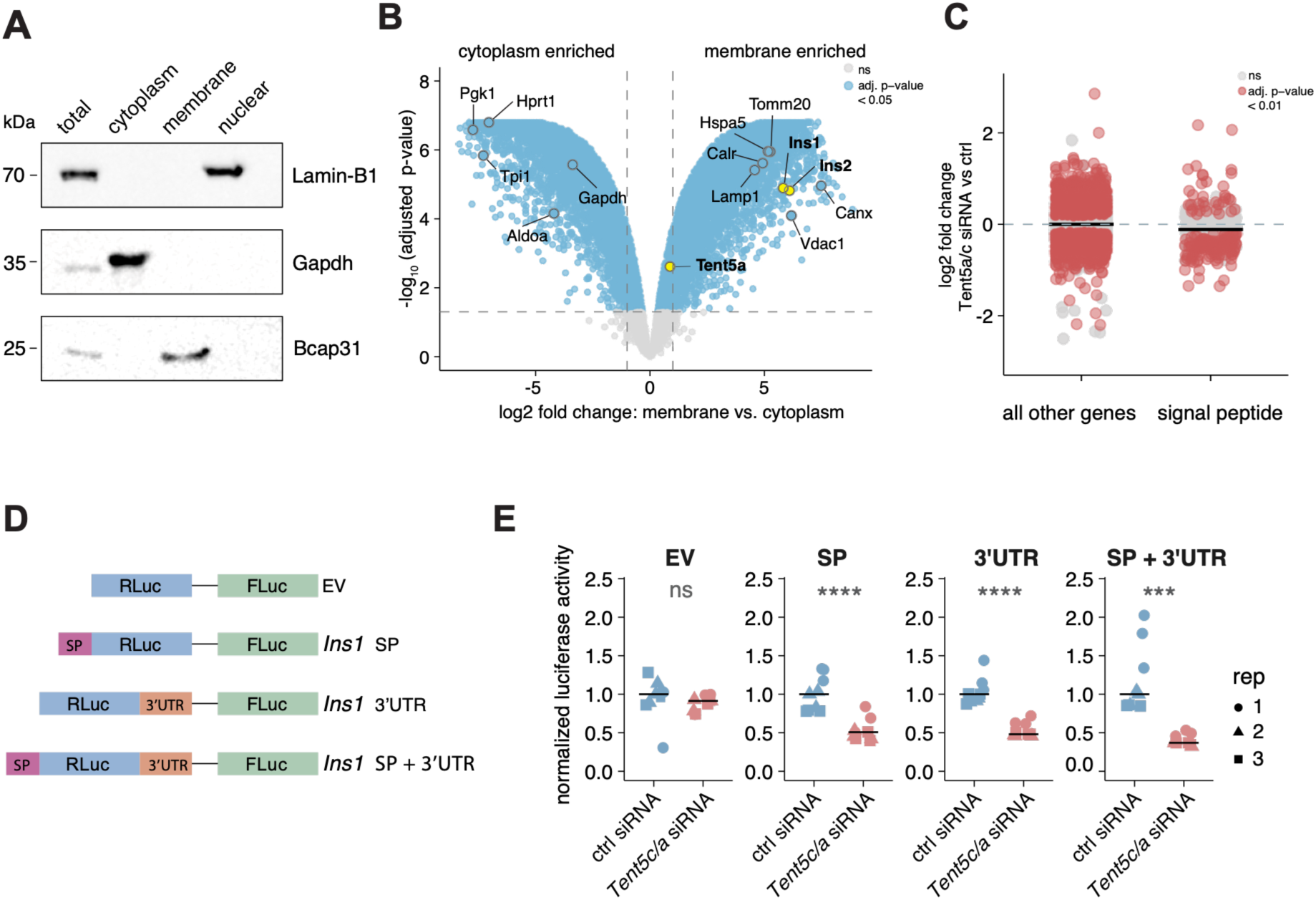
Insulin mRNA regulation through Tent5 is stimulated by both its 3’UTR and ER localization. A. Subcellular fractionation in INS1-E cells. Representative Western Blot analysis measuring nuclear (Lamin-B1), cytosolic (Gapdh) and endoplasmic reticulum (Bcap31) markers in subcellular fractions and whole extracts (n=3 independent replicates, additional two replicates in Suppl. Fig.4B) B. Subcellular localization of Tent5a assessed by quantitative proteomics. Label-free LC–MS/MS analysis comparing membrane and cytosolic fractions isolated from INS-1E cells. Each point represents a detected protein, plotted according to its log2fold change (membrane to cytosolic) and −log₁₀ adjusted p-value. Enrichment of canonical cytosolic (Gapdh, Pgk1, Hprt1, Aldoa) and membrane-associated (Ins1, Ins2, Canx, Calr, Lamp1, Tomm20, Vdac1, Hspa5) markers demonstrates the purity of the respective fractions. C. Distribution of log2 fold changes (*Tent5a/c* siRNA vs ctrl siRNA) stratified by the presence of a predicted signal peptide (SP) as annotated for rat genes (SignalP6, SEPDB open(111)) (n=3, independent experiments). Genes with adjusted p-value < 0.01 are highlighted in red, while non-significant genes are shown in grey. Crossbars indicate medians for each group. D. Schematic representation of luciferase reporters containing no fusion (EV), the rat Insulin 1 (*Ins1*) signal peptide (SP) upstream; the 3’UTR downstream the *Renilla luciferase* coding sequence or both. E. Normalized *Renilla luciferase* activities of reporters carrying no fusion (EV), the rat Insulin 1 (*Ins1*) signal peptide (SP), its 3′UTR, or both, in control INS-1E cells (ctrl siRNA) or *Tent5a/c* double knockdown cells (*Tent5a/c* siRNA) (n = 3 independent experiments, each with technical triplicates). *Renilla luciferase* activities were first internally normalized to firefly luciferase activity and then expressed as fold-change relative to the corresponding control condition; crossbars indicate medians. Paired t-test p-values indicated as follows: <0.001 (***), <0.0001(****).

To test whether the specificity of Tent5 activity on insulin mRNAs is linked to their ER localization, we cloned the *Ins1* signal peptide (SP) in fusion with a luciferase reporter (Fig. 4D) and combined a reporter assay with *Tent5a/c* knock-down in INS-1E cells. We observed that the signal peptide reduces luciferase activity in control conditions, as previously reported (68,69) (Suppl. Fig. 4F). *Tent5a/c* knock-down resulted in a 40% reduction of the luciferase activity in presence of the insulin SP, while the reporter alone showed substantially unperturbed luciferase activity (Fig. 4E). Conversely, we also cloned the *Ins1* 3’UTR downstream of the reporter coding sequence and observed that *Tent5a/c* knock-down resulted in a reduction in luciferase activity comparable to that obtained with the SP alone (Fig. 4E). Finally, the loss in luciferase activity upon *Tent5a/c* knock-down was slightly stronger in the presence of both the insulin SP and its 3’ UTR (Fig. 4E). Of note, the presence of the insulin 3’UTR was sufficient to enhance luciferase activity by ca. 50% both in control conditions and in the presence of the insulin SP (Suppl.Fig.4F).

From these results, we conclude that the ER localization of Tent5 enzymes accounts for only a part of their substrate specificity, and we show that the insulin 3’ UTR is also sufficient to stimulate Tent5-mediated stabilization of a reporter mRNA.

### 5. Insulin deadenylation and Tent5a/c abundance are dynamically regulated by glucose

Given the notion that insulin mRNA metabolism is regulated by glucose (2–27) and having established the role of Tent5-in regulating insulin mRNA in basal conditions, we tested whether this mechanism is itself dynamically regulated. We subjected INS-1E cells to a 1-hour pulse of glucose deprivation followed by 3 hours of high glucose stimulation and measured genome-wide poly(A) tail lengths. Strikingly, we observed a strong shortening of both *Ins1* and *Ins2* poly(A) tails upon acute glucose stimulation (Fig. 5A, Suppl. Fig 5A). In addition, we observed glucose-induced poly(A) tail shortening also in highly abundant hormone peptides and factors involved in granule biogenesis and which were significantly downregulated in *Tent5a/c* knockdown (Fig.3B), such as *Chga/b* and *Iapp* (Suppl. Fig. 5B)

**5.**
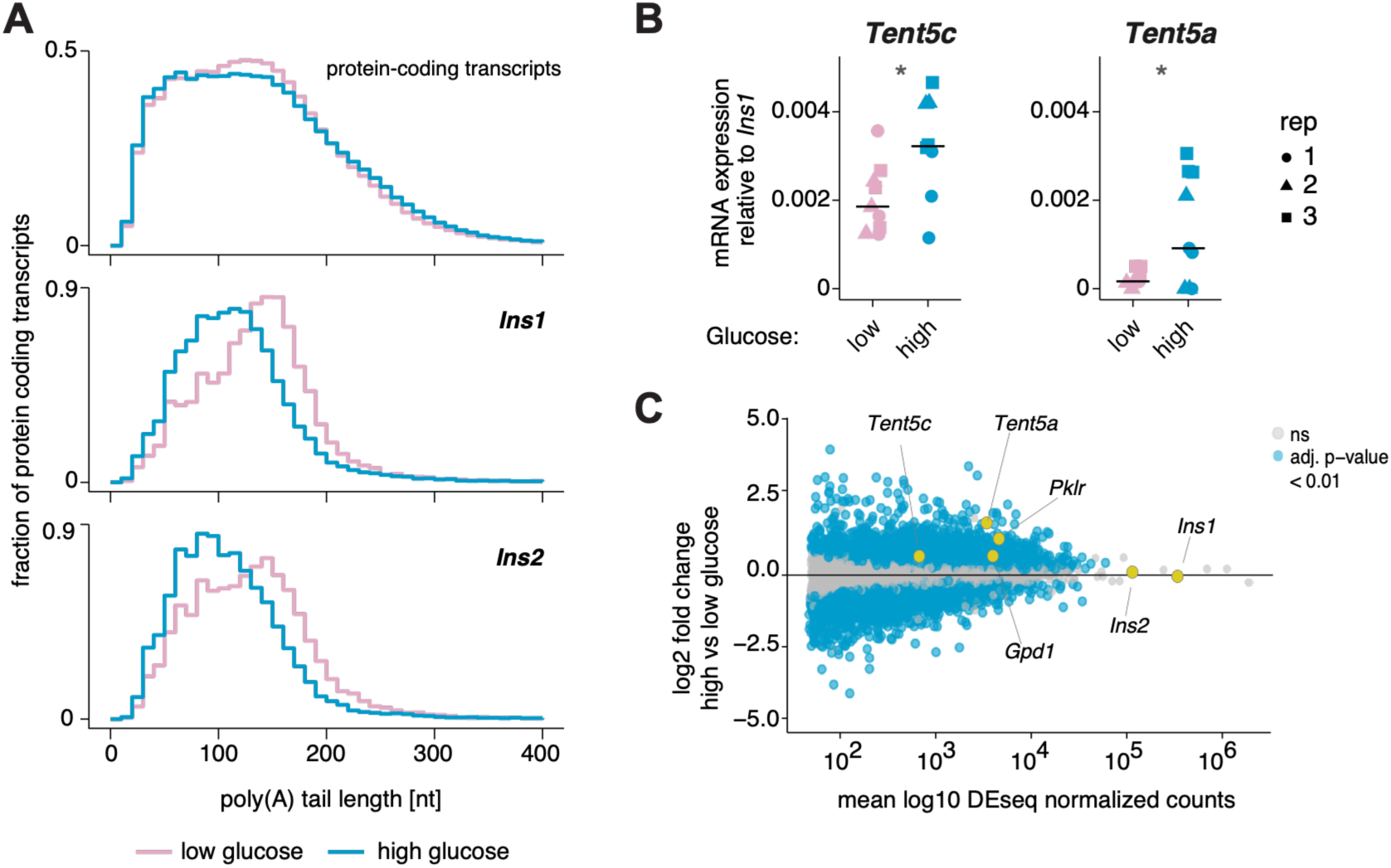
Insulin poly(A) tail shortening and *Tent5a/c* expression are induced by glucose. A. Poly(A) tail length distributions for Ins1/2 and all protein coding transcripts upon 1 hour glucose starvation (low glucose, 2.8mM) or 3 hours acute glucose stimulation (high glucose, 28mM) in INS-1E cells. Aggregated results of n=2 independent experiments. Distributions are normalized so that the area under the curve equals 1 are shown. B. qRT-PCR analysis of *Tent5c* and *Tent5a* in INS-1E cells equilibrated in low glucose media (2.8mM, 1 hour) or stimulated with high glucose media (28mM, 3 hours). Expression was normalized to Ins1, which remains stable upon 3 hours acute glucose stimulation (see Fig.3C). Data represent n = 3 independent experiments, each with technical triplicates; crossbars indicate medians. Paired t-test p-values indicated as follows: <0.05 (*). C. MA plot of differential gene expression comparing control siRNA-treated INS-1E cells in high (28mM) vs low (2.8mM) glucose conditions (n=3, independent experiments). Differential expression is plotted as the log2 fold change (high vs low glucose) versus the mean DESeq2-normalized expression. Genes with adjusted *p* < 0.01 are highlighted in blue; non-significant genes are shown in grey. Selected genes are annotated in outlined or yellow circles. Differential expression was assessed using DESeq2; genes with mean normalized counts < 50 were filtered prior to testing.

We then measured *Tent5a/c* mRNAs abundance by qRT-PCR (Fig. 5B) and bulk RNA-seq (Fig. 5C). In agreement with previous reports (e.g. 70), glucose stimulation exerted a dramatic effect on gene expression, and we observed both *Tent5a* and *c* being rapidly upregulated to an extent similar to that of some well-known targets of the glucose sensing transcription factor ChREBP, such as *Gpd1* and *Pklr* (Fig. 5C). Of note, the observed transcriptional changes were only mildly affected by *Tent5a/c* knock-down (Suppl. Fig. 5C), indicating that their activity plays a role downstream of the glucose response. In fact, *Ins1/2* were still strongly downregulated right after starvation and after glucose stimulation (Suppl. Fig. 5D-E).

Overall, our observations indicate that the insulin mRNA is rapidly deadenylated upon acute glucose stimulation, while the *Tent5a/c* genes are likely transcriptionally activated, possibly to restore the insulin mRNA pool which would be otherwise depleted.

## Discussion

### Insulin mRNA metabolism is tightly regulated in beta cells

Insulin is a key metabolic hormone controlling glucose homeostasis in vertebrates, and its dysfunction results in diabetes with severe health complications. It is therefore essential to understand the key principles underlying the regulation of insulin biosynthesis and activity. While high glucose levels result in the fast release of insulin in preformed secretory granules by beta cells, insulin biosynthesis is also tightly regulated at the mRNA level, although in a highly complex, stimulus- and time-dependent manner. Here we report direct evidence that short-term insulin mRNA stabilization upon glucose stimulation can be at least partially attributed to the activity of Tent5a and c poly(A) polymerases, which are rapidly induced upon glucose stimulation.

Consistent with this dynamic regulation of insulin mRNA, transcription of either the *INS* gene in humans or the two non-allelic *Ins1* and *Ins2* genes in rodents has been shown to be induced by short- or medium-term glucose stimulation (8–11) and reduced upon chronic glucose exposure (12–14). Moreover, it has been reported that the insulin mRNA is rapidly sequestered into granules upon starvation, and those granules dissolve upon acute glucose stimulation to make the insulin mRNA again available for translation (71). Preproinsulin translation has been in fact shown to be induced by glucose through multiple mechanisms involving the activity of proteins binding their 5’ UTR (15–21), but reduced by chronic glucose exposure prior to or as a consequence of ER stress, mainly via EIF2alpha phosphorylation (22–24,72,73). Finally, insulin mRNA decay was shown to be triggered by prolonged high glucose exposure and ER stress, due to the activation of the UPR (25,74), and particularly the IRE1alpha nuclease (75,76), although its stability was found to be enhanced in response to short glucose stimulation (26,27).

### A mechanism involving insulin regulation by TENT5 poly(A) polymerases

Both translation and stability of mRNA depend on the poly (A) tail: after export into the cytoplasm, translation of polyadenylated mRNAs is stimulated by poly(A) binding proteins (77), while at the same time the tail is shortened by the action of deadenylases (78,79) at a gene-specific rate (31–34,80,81), until it reaches a critical length below which mRNA undergoes irreversible exonucleolytic decay (82). Re-adenylation in the cytoplasm can therefore stabilize mRNAs, counteracting the action of deadenylases. The TENT5 family proteins have been shown to selectively stabilize mRNAs encoding for secreted proteins or proteins involved in folding, glycosylation and trafficking via re-adenylation (38). TENT5 proteins localize at the ER via the interaction with FNDC3 proteins (49) and regulate several biological processes, such as proliferation and antibody production in B cells (39,50), gametogenesis (53,83), osteogenesis (84) and erythropoiesis (85).

Here we found that the *TENT5* family genes *TENT5A and C* are particularly highly expressed in the human pancreas, and within pancreas *TENT5C* is most abundant in endocrine cells including beta cells, while *TENT5A* is more ubiquitously expressed. We found that *TENT5C* is genetically associated with glycemia and T2D, which was also reported and validated independently in a study aimed at identifying the genetic basis of beta cell function (86), although these findings do not necessarily implicate a direct role in beta cells. Importantly, *TENT5C* was also among the hits of a CRISPR screen aimed at identifying beta cell function modifiers (87). Given the previously reported role of TENT5 proteins in stabilizing ER-targeted mRNAs, we hypothesized that they might be directly involved in insulin mRNA regulation. By compiling an atlas of genome-wide poly(A) tail length distributions in the human pancreas, we observed that a number of mRNAs encoding for abundant peptides produced by endocrine cells, including the insulin mRNA, have a particularly long poly(A) tail compared to most mRNAs expressed in the same tissue. We therefore performed knock-down experiments for *Tent5a* and *c* in rat INS-1E cells and found that their depletion results in a strong, significant reduction in the abundance of the insulin mRNA, as well other hormone peptide-encoding mRNAs and mRNAs encoding for factors involved in secretory granules biogenesis (e.g. *Chga/b*). To prove that this effect was directly linked to *Tent5a/c* readenylation activity, we measured poly(A) tail length distributions upon *Tent5a/c* depletion, but we only observed a mild effect for the cytosolic pool of *Ins1/2* mRNAs. While we were finalizing this work, Silva, Mayrhofer et al. published very similar findings (67) and suggested that rapid decay of deadenylated mRNA likely masks the effect on tail length distributions at steady state. In fact, they observed that overexpression of Tent5a causes an increase in insulin mRNA tail length, confirming it as a substrate for readenylation. While we agree that this interpretation is sensible, we note that we did observe stronger effects on steady state tail lengths for other Tent5a/c-regulated mRNAs, indicating that additional or alternative regulatory mechanisms might explain the observations for the insulin mRNA pool.

So far, the only described mechanism for Tent5 substrate specificity is based on subcellular localization, involving ER-residing mRNAs being preferentially re-adenylated by ER-docked Tent5 proteins. We confirmed that Tent5a is enriched at the ER in a beta cell line, and that the insulin signal peptide is sufficient to elicit Tent5a/c-mediated stabilization of a reporter mRNA. However, we observed that signal peptide-encoding mRNAs do not tend to carry longer poly(A) tails in TENT5-expressing tissues, nor that these mRNAs globally undergo destabilization or deadenylation upon Tent5 depletion. Moreover, we found that the insulin 3’ UTR alone can stabilize a reporter mRNA and responds to Tent5 activity to a similar if not to a larger extent than the insulin signal peptide. These findings suggest that either the insulin 3’ UTR itself acts as an ER localization signal, and/or additional specificity factors possibly facilitate the interaction of Tent5 with its substrates. Given the recent observation that Tent5-mediated re-adenylation enhances the efficacy of mRNA vaccines (40), we propose that identifying these specificity factors and their target cis-regulatory elements may directly impact on RNA therapeutics applications. To this regard, we anticipate the potential use of the insulin 3’UTR or modified, synthetic alternatives aimed at stabilizing mRNAs via re-adenylation and/or targeting them to the secretory pathway.

Finally, our observation that glucose stimulation induces rapid poly(A) tail shortening of insulin mRNA provides a mechanistic explanation for the need of re-adenylation, as repeated glucose exposures would rapidly deplete beta cells of their insulin mRNA pool. Consistently, *Tent5a/c* seem to be transcriptionally activated upon glucose stimulation, thus providing a compensatory mechanism for glucose-dependent tail shortening.

### A potential role for TENT5 activity in the pathogenesis of T2D?

In T2D, beta cells compensate insulin resistance by increasing preproinsulin synthesis, which burdens ER homeostasis causing ER stress and ultimately, if unresolved, cell death, (42). The Unfolded Protein Response (UPR) has in fact been shown to play an essential role in protecting beta cells from ER stress (24,72,88–91), for example via IRE1alpha-mediated degradation of the insulin mRNA (25,74,75). Of note, misfolding of proinsulin at the ER of beta cells is an early molecular hallmark of the progression from prediabetes to overt diabetes in mice (42). Mounting evidence therefore proposes ER stress as a pathogenetic mechanism for T2D, and its targeting as a promising novel strategy for early interception (92,93). Along this line, it has been shown that reducing insulin synthesis in mice alleviates ER stress and improves beta cell function, and in general protects against obesity and diabetes (46–48). Consistently, it was shown that targeting key effectors of the UPR (45,94), as well as the use of GLP1 analogues (43,44) can relieve ER stress and improve beta cell function. Within this context, it is important to note that TENT5C was shown to boost ER growth and secretory output in multiple myeloma cells beyond their capacity, causing ER stress and apoptosis (49).Here we further report that variants in the *TENT5C* gene are strongly associated with T2D and glycaemia, and that Tent5a and *c* expression is regulated by glucose in rat cells. Our findings demonstrate that TENT5 proteins sustain insulin biosynthesis in physiological conditions, an activity that may sensitize beta cells to ER stress in pathological conditions. Based on this model, we propose that *TENT5A/C* may be explored as potential targets within the context of the emerging therapeutic opportunities aimed at the early interception of T2D by targeting ER stress.

### Limitations of this study

Overall, here we use genome-wide profiling of poly(A) tail length, single-cell and bulk transcriptomics, genetic evidence and perturbation experiments in *in vitro* beta cell models to report a novel mechanism sustaining insulin biosynthesis in physiological conditions, which is linked to T2D pathogenesis.

While the genetic association of *TENT5C* with T2D and its perturbed expression in islets from T2D patients come directly from human data, our findings that TENT5 enzymes control insulin mRNA metabolism rely on *in vitro* models and will need to be confirmed in more physiological *ex vivo* islet explants or *in vivo*. To this regard, we note that mild glucose intolerance was reported in *Tent5c* knock-out mice (86,95) and that TENT5A overexpression increased insulin content in human islet microtissues (67).

Furthermore, we cannot at present disentangle the relative contribution of TENT5A and C to this mechanism in human cells. While *Tent5a* is more abundant than *Tent5c* in rat INS-1E cells, both seem to exert an effect on insulin mRNA. In human islets, we observe *TENT5C* being more abundant at the RNA level, while TENT5A seems more abundant in beta cells although this information relies on single-cell RNA-seq alone which is more prone to gene-to-gene variability artefacts. Therefore, whether these two proteins are redundant remains to be tested in human beta cell models. Moreover, further investigation is necessary to determine the exact mechanism ensuring TENT5 target specificity and poly(A) tail length control, as well as the exact nature of the interplay between TENT5 activity and ER stress. Along this line, the hypothesis that TENT5-mediated stabilization of the insulin mRNA may be linked to T2D pathogenesis and therefore amenable to potential therapeutic interventions remains to be explored, especially in the light of the multiple roles of TENT5 proteins in the physiology of many other secretory cell types and their role as an oncosuppressor (49).

## Acknowledgements

We thank Oliver Harschnitz, Monika Piwecka, Agnieszka Rybak-Wolf, Shruti Bhagat, Yasuhiro Murakawa, Enrico Milan, Antonio Astorino and Manendra Singh Negi for the precious discussions and feedback. We acknowledge the access and services provided by the National Facility for genomics (IU1 – High-throughput sequencing), genome engineering and disease modelling (IU2 – Gene editing technologies), light imaging (IU1 – Imaging), data handling and analysis (IU1 – Bioimage analysis), structural biology (IU4 – Structural proteomics), and particularly thank Chiara Ambrosini, Eugenia Ricciardelli, Alessandra Fasciani, Eugenia Cammarota and Andrea Graziadei. We thank Iren Xhaferi for support with illustrations. We used AI (ChatGPT v. 5.2 and M365 Copilot) to help with code writing/editing. Ana Kotte and Lucia Prandi are PhD students within the European School of Molecular Medicine (SEMM).

## Author contributions

AK, FM, NvW, LM, LP, SB, JZ and AZ performed experiments. AK, FM, NvW, LM, LP, AZ, RP, RSS, IL and BS performed analyses. IL and AZ supervised the project.

## Data and code availability

This manuscript used publicly available data from EBI (E-MTAB-5060 and E-MTAB-5061), GEO (GSE153855, GSE214517), the database (https://hpap.pmacs.upenn.edu/) of the Human Pancreas Analysis Program (HPAP; RRID:SCR_016202)(62,63). HPAP is part of a Human Islet Research Network (RRID:SCR_014393) consortium (UC4-DK112217, U01-DK123594, UC4-DK112232, and U01-DK123716). Raw sequencing data generated in this work are deposited on SRA under accession number PRJNA1484008. Human sequencing data are available upon data transfer agreement. Mass spectrometry data have been deposited to the ProteomeXchange Consortium via the PRIDE partner repository with the dataset identifier PXD079616. The pipeline used to process FLAM-seq data is available under https://github.com/Legnini-Group/SnakeFLAME/.

## Materials and Methods

### Cell culture

We cultured INS-1E Rat insulinoma beta cells (Sigma, cat. #SCC491) in T25 flasks (Sigma-Aldrich, cat #SCC491) in RPMI-1640 with L-Glutamine (Euroclone, cat #ECB2000L) supplemented with 10% Fetal Bovine Serum (Euroclone, cat #ECS5000LH), 10 mM HEPES (Fisher bioreagents, cat. #BP310-1), 1 mM Sodium Pyruvate (Sigma-Aldrich, cat. #S8636), and 50 µM 2-Mercaptoethanol (Gibco, cat. #21985023). We maintained cells at 37°C in a humidified incubator with 5% CO_2_ and passaged them once per week. For passaging, we rinsed cells with 5 mL PBS, detached them using 1 mL Accutase (Euroclone, cat #ECB3056D) for 5 minutes at 37°C, and centrifuged them at 300 × g for 2-3 minutes. We gently resuspended the cell pellet in complete growth medium and plated the cells as required. We used cells between passages 20 and 40 in all experiments.

### Transfections

We seeded INS-1E cells into 6-well plates and maintained them at 37°C in a humidified incubator for 72 hours to ensure complete adherence. For *Tent5a/c* knockdowns, we incubated a transfection mixture containing either scrambled control siRNA (siTOOLs, cat#SI-C005) or a combination of *Tent5a* siRNA (siTOOLs, cat #si-G050-300870) and *Tent5c* siRNA (siTOOLs, cat #si-G050-310721) with Lipofectamine 3000 (without P3000 reagents) in Opti-MEM (Gibco, cat #31985062) at room temperature for 15 minutes. We replaced the culture medium with fresh complete growth medium and added the transfection mixture dropwise directly to the cells to achieve final concentrations of 48 nM for the scrambled control siRNA or 24 nM for each *Tent5a* and *Tent5c* siRNA. We then incubated cells at 37°C for 48 hours before downstream processing.

### Glucose Stimulation

We seeded and transfected INS-1E cells as described above. Forty-eight hours following transfection, we equilibrated the cells in a modified KREBS buffer containing 25 mM HEPES, 115 mM NaCl, 5 mM KCl, 1 mM MgCl_2_, 2.5 mM CaCl_2_, 0.2% (w/v) BSA supplemented with 2.8 mM glucose (Thermo Fischer Scientific, cat #A2494001) for 1 hour. We collected conditioned medium after 1 hour for ELISA analysis, and harvested RNA at this stage from a parallel plate. We then stimulated the cells by exposing them to high-glucose (28 mM) KREBS buffer. After 3 hours of stimulation, we collected the supernatant for ELISA, and lysed cells in TRI reagent for downstream RNA extraction (Zymo Research, cat #R2052).

### Plasmids

We cloned the signal peptide of the rat *Ins1* gene upstream and in-frame with the Renilla luciferase coding sequence in the dual-luciferase reporter vector psiCheck2 (Promega, cat. #C8021). For 3′ UTR reporter constructs, we designed an insert to encompass the Ins1 3′UTR, the canonical polyadenylation signal (PAS, AATAAA), followed by 41 nucleotides of the genomic sequence downstream of the PAS. We cloned the insert into XhoI-NotI restrictions sites downstream of the Renilla luciferase coding sequence in either the unmodified psiCheck2 backbone or the modified backbone containing the Ins1 signal peptide. The exact cloned sequences are available in Suppl. Table 1.

### Cell fractionation

We passaged INS-1E replicates independently, as described above, until they reached ∼80% confluency (∼2 × 10⁷ cells per replicate) and subjected them to sequential detergent-based subcellular fractionation as previously described(96). We washed the cells once with PBS, detached them using Accutase (Euroclone, cat. #ECB3056D) for 5 minutes at 37 °C, collected the cell suspension in 5 mL ice-cold PBS supplemented with 100 μg/ml cycloheximide (Sigma-Aldrich cat. #C4859), and pelleted the cells by centrifugation for 4 min at 300 x g.

We resuspended the cell pellets in 400 µL ice-cold permeabilization buffer (110 mM KOAc, 25 mM K-HEPES pH 7.2, 2.5 mM Mg(OAc)₂, 1 mM EGTA) supplemented with 0.015% v/v digitonin, 1 mM DTT, 100 μg/ml cycloheximide (Sigma-Aldrich cat. #C4859) and 1x EDTA-free protease inhibitor cocktail (Roche cat. #11836170001), incubated at 4 °C with rotation for 10 minutes, and centrifuged at 3000 × g for 5 minutes at 4 °C. We collected the supernatant as the cytoplasmic fraction. We then washed the pellets twice in 1 mL ice-cold washing buffer (permeabilization buffer containing 0.004% digitonin), followed by centrifugation at 3000 × g for 5 minutes at 4 °C after each wash. We lysed the washed pellets in 400 µL ice-cold lysis buffer (400 mM KOAc, 25 mM K-HEPES pH 7.2, 15 mM Mg(OAc)₂, 0.5% v/v NP-40, 1 mM DTT, 100 μg/ml cycloheximide (Sigma-Aldrich cat. #C4859), 1X-EDTA free protease inhibitor cocktail (Roche cat. #11836170001), incubated the samples on ice for 5 min, and centrifuged them at 3000 × g for 5 minutes at 4 °C. We collected the supernatant as the membrane-associated fraction, and the pellets the nuclear insoluble fraction. For further purification, we loaded the nuclear pellet onto a 10% sucrose cushion in lysis buffer and centrifuged at 200 × g for 5 minutes at 4 °C; this step was repeated once. We resuspended the resulting nuclear pellet in 400 µL 1× RIPA buffer (50mM Tris-HCl pH 7.4, 150 mM NaCl, 0.5% v/v sodium deoxycholate, 1 % v/v Triton X-100, 0.1% v/v SDS, supplemented with 1x EDTA-free protease inhibitor cocktail (Roche cat. #11836170001) and incubated on ice for 5 minutes. We prepared total protein lysates in parallel by directly lysing cell pellets in 1× RIPA buffer for 10 minutes on ice.

We clarified all fractions (total, cytoplasmic, membrane, and nuclear) from cell debris by centrifugation at 7500 × g for 10 minutes at 4 °C and collected aliquots (20 µL) for western blot analysis. The remaining fractions were mixed with 3x volumes Trizol LS (Thermo Fisher cat. #10296010) for subsequent RNA extraction.

### Western Blot

We performed Western Blot analysis according to standard protocols(97). Briefly, we resolved 20 µL protein lysate on a 10% SDS PAGE, transferred to a nitrocellulose membrane (Cytiva cat. #15259794) and blocked in blocking buffer (5% skim-milk in TBST; 1x Tris-buffered saline, 0.1% Tween-20). For cellular fractionations, we probed membranes with primary antibodies against a cytosolic marker (GAPDH, Proteintech cat. #60004-1-Ig), ER marker (BCAP31, Proteintech, cat. #11200-1-AP), nuclear marker (LAMIN-B1, Abcam cat. #AB16048) diluted according to manufacturer’s recommendations in blocking buffer. We washed the membranes three times in 1x TBST (1x Tris-buffered saline, 0.1% Tween-20) and probed them with secondary antibodies for 1h at room Temperature, anti-mouse-HRP (Invitrogen cat. #SA1-100) or anti-rabbit-HRP (Invitrogen cat. #31458), diluted 1:10,000 blocking buffer. Following three additional washes in TBST, we probed membranes with Pierce ECL Western Blotting Substrate (Thermo Fisher Scientific cat. #32106) and imaged on an Odyssey M imaging system (LICORbio).

### Luciferase assays

For luciferase assays: we seeded INS-1E at a density of 50,000 cells per well in 200 µL medium on white-walled 96-well plates, cultured to 60-70% confluency and co-transfected with siRNAs (48nM negative control siRNA; siTOOLs cat. #SI-C005, or a combination of 24nM *Tent5c* siRNA; siTOOLs cat. #si-G050-310721, and 24nM *Tent5a* siRNA; siTOOLS cat. #SI-G050-300870) and a luciferase reporter (0.5 µg/mL), either the psiCheck2 backbone or one of the modified reporter constructs. Forty-eight hours post-transfection we assayed firefly and renilla luciferase activities using the Dual-Glo Luciferase Assay System (Promega cat. #E2920) and measured luminescence on a GloMax Discover microplate reader. We normalized Renilla luciferase activity to firefly luciferase activity internally, following background subtraction of luminescence from non-transfected cells.

### Immunostainings

For INS-1E, we seeded cells at a density of 40,000 cells per well in 200 µL final volume of medium on a 96-well plate (Ibidi cat. #89626). Three days after seeding we washed cells three times in 1X PBS, fixed them by incubation in 4% PFA (Thermo Fisher Scientific cat. #10553151) for 15 minutes at room temperature, followed by three additional washes in 1X PBS. We performed permeabilization in 1X PBS containing 0.5% Triton X-100 (Sigma-Aldrich, cat. #9036-19-5) for 15 minutes, followed by three washes in 1X PBS and subsequent blocking by incubating in blocking solution (5% donkey serum, 0.1% Tween-20 in 1X PBS) for 45 minutes at room temperature. We incubated the cells overnight at 4°C with three different primary antibodies against TENT5C (Bioss Antibodies, cat # bs-8198R, Novus Biologicals, cat # NBP3-48488, Invitrogen, cat #PA5-119549) diluted 1:50, 1:100 and 1:500 respectively in blocking solution. After three washes with 1X PBS, we incubated cells with a mixture containing secondary antibody (Life Technology, cat. #A-10042) diluted 1:500 and phalloidin diluted 1:400 in blocking solution for 1 hour at room temperature protected from light. Next, we washed cells thrice with 1X PBS and performed nuclear staining using DAPI solution (10 mg/mL) diluted 1:5000 in blocking solution for 10 minutes at room temperature. We then washed the cells three additional times with 1X PBS. We acquired images with an inverted Leica Stellaris 8 confocal microscope equipped with a Plan Apochromatic 63x/1.4 Oil CS2 objective. We used the integrated white light laser (WLL), with excitation wavelengths selected for DAPI (405 nm), TENT5C (secondary: ALEXA 555 nm), and Phalloidin (ALEXA 647 nm). We detected Emission signals using Leica HyD X and HyD S detectors. We acquired images in 16 bit and kept laser power and detector gain constant across comparable samples. Microscopy acquisitions were performed at the National Facility for Light Imaging (IU1-Imaging), Fondazione Human Technopole, Milan, Italy.

### Imaging and single-cell intensity analysis

We performed colocalization analysis using ImageJ (Fiji). Firstly, we split the image into the four channels acquired. We used Channels 1 (DAPI) and 2 (phalloidin) as input for Cellpose (v4.0.7) to perform cell segmentation. We used Regions of interest (ROIs) retrieved from cell segmentation for downstream analysis. We first computed the mean intensity for each segmented cell within each field of view (FOV) acquired. We then calculated the median of all single-cell ROI mean intensities within each FOV and used this value for statistical comparisons between conditions.

### Liquid chromatography/mass spectrometry (LC-MS) based proteomics

We fractionated 2 × 10⁷ INS1-E cells in biological triplicates to obtain cellular compartments. We processed them by single-pot, solid phase proteomics sample preparation (SP^3^) on an Agilent Bravo LT96 liquid handling platform (98) using Sera-mag beads (Cytiva cat # 44152105050250 and 24152105050250 mixed 1:1). We used 7 micrograms of proteins with a 10:1 w/w ratio of beads to proteins and 18 minutes binding time. We performed reduction with 2.5 mM dithiothreitol for 30 minutes at 37 °C, followed by alkylation with 5 mM iodoacedamide for 30 minutes at 37 °C. The proteins were then digested with trypsin overnight at 37 °C.

We injected peptides for each liquid chromatography-mass spectrometry (LC-MS) acquisition. The LC-MS platform consisted of a Vanquish Neo system (Thermo Fisher Scientific) connected to an Orbitrap Astral mass spectrometer (Thermo Fisher Scientific) operating under Tune 2.0.690.17. Mobile phases consisted of 0.1% v/v formic acid in water (mobile phase A) and 0.1% formic acid in 80% acetonitrile/water v/v (mobile phase B). We set up the system in trap-and-elute workflow with a pepmap Neo trap cartridge (Thermo Fisher Scientific cat # 174500). We acquired 40 samples per day and separated on an EASY-Spray PepMap Neo column (150 µm x 15 cm) (Thermo Fisher Scientific cat # ES906) with a 33-minutes gradient employing a linear separation between 8%-22.5% B at 800 nl/min in 24 minutes.

We performed MS acquisitions in narrow-window data-independent (DIA) mode (99). We acquired MS1 spectra with a resolution of 240000 and 5 ms maximum injection time with a full scan range of 380–980 m/z and automatic gain control of 500%, setting Source RF lens to 40%. We performed the DIA part with 300 windows of 2-Th scanning from 380 to 980 m/z under a time control of 0.6 second maximum second per duty cycle, setting the ion injection time to a maximum of 3ms per windows and the automatic gain control target to 500% (50000 ions). The isolated ions were fragmented using HCD with 25% Normalized Collision Energy.

We processed the resulting raw files with DIA-NN 2.3.0(100) using an *in silico* spectral library generated by DIA-NN from *R. norvegicus* UniProt (including TrEMBL, all isoforms and common MS contaminants). The spectral library is included in the data deposition. The spectral library included methionine oxidation, 1 missed cleavage, precursor charge states 1-4 and common contaminants. We searched the data with a minimum peptide length of 7, 1 missed cleavage, methionine oxidation as variable modification, cysteine carbamidomethylation as fixed modification, match between runs, and the direct quantification option in DIA-NN. We disabled normalization and set proteotypicity to genes. We filtered the data with a 0.01 q-value filter at the precursor level and 1% false discovery rate (FDR) at the protein group level.

We analyzed the unique gene matrix (report.unique_genes_matrix.tsv) using QProMS (https://github.com/ieoresearch/QProMS), selecting proteins present in a minimum of 80% of replicates in at least one condition. This resulted in an identification of 10,031 protein groups for the whole cell proteomes. We normalized the values with variance-stabilizing normalization (VSN) (101) and carried out imputation with the missForest algorithm (102,103). We carried out univariate enrichment analysis with the Welch test with Benjamini-Hochberg FDR correction for multiple testing (alpha=0.05). Mass spectrometry was performed at the National Facility for Structural Biology (IU4 – Structural proteomics), Fondazione Human Technopole, Milan, Italy.

### ELISA

We collected the supernatant of INS-1E cells from the glucose stimulation assay described above. To measure insulin content in the media, we performed ELISA using the Mercodia Ultrasensitive Rat Insulin kit (Mercodia cat. #10-1251-01), following the standard protocol. To ensure optimal range of concentration, we diluted all the samples 1:1000 or 1:500 in Calibrator 0 provided with the kit. We used the VICTOR Nivo Multimode Reader (PerkinElmer, HH3500) to read the absorbance at 450 nm and used these data to derive insulin concentration in each sample.

### RNA purification and qRT-PCR

To extract RNA we used the Mini-prep Direct-Zol RNA purification kit (Zymo Research cat. #R2052) including a DNase I digestion step. RNA from INS-1E subcellular fractions was purified using standard phenol-chloroform extraction including a Turbo DNase (Thermo Fisher, cat. # AM2238) digestion step according to the manufacturer’s recommendation. We carried out reverse transcription with 20-100 ng of RNA using 50 ng random primers (Promega cat. #C1181) and Superscript IV (Invitrogen, cat. #18090200) according to the manufacturer’s protocol. We then performed real time qPCR in 10 µL reactions using 1 µL resulting cDNA, PowerUp SYBR Green Master Mix (applied biosystems cat. #A25776) and 0.5 µM forward and reverse primers (listed in Suppl. table 1). We ran samples in triplicate on an applied biosystems QuantStudio 5 Real-Time PCR system (Thermo Fisher Scientific) under the following conditions: 2min at 50°C, 10min at 95°C, 40 cycles of 15s at 95°C and 1min at 60°C and a final melt curve stage using the QuantStudio Design and Analysis Software (v1.5.3). Within each sample, we normalised gene expression to the chosen reference gene using 2^-ΔCt^ to calculate the relative expression.

### Bulk RNA-seq

For bulk RNA-seq, we prepared libraries using the Illumina Stranded Total RNA Prep, Ligation with Ribo-Zero Plus kit (Illumina, cat. #20040529). We amplified the libraries via 14 cycles of PCR and sequenced on Illumina NovaSeq 6000, S2 flowcell, PE 100 (Illumina, cat. #20028315), generating an average number of 45 million reads per sample. Bulk RNA-seq was performed at the National Facility for genomics (IU1 – High-throughput sequencing), Fondazione Human Technopole, Milan, Italy.

For the re-analysis of publicly available RNA-seq data, we ran Nextflow v. 25.10.3(104) on FASTQ files (s. Acknowledgements and Data availability sections) with standard parameters in a container using Singularity v. 3.8.5, using the UCSC human GRCh38 genome and GRCH38 v39 Gencode annotation. We then used raw count tables for downstream differential gene expression analyses with Deseq 2 v. 1.46.0(105) in RStudioPro v. 2024.04.2 and R v. 4.4.0, and TPMs calculated with the Nextflow Star/Salmon(106,107) quantification pipeline for all other purposes. For the analysis of in-house-produced bulk RNA-seq data, we used a similar, custom Nextflow-based bulk RNA-seq pipeline made available by the National Facility for Genomics at Human Technopole.

### Single-cell RNA-seq

We aligned raw FASTQ files from the human pancreatic islet single-cell RNA-seq dataset E-MTAB-5061(58) and GSE214517(59) to the GRCh38 reference genome using STAR (v. 2.7.9a). We generated two different indexed genomes accounting for the differences in read length of the two datasets using STAR, setting the --sjdbOverhang parameter to read length – 1. We ran STARsolo in SmartSeq mode (--soloType SmartSeq) to generate single-cell gene expression count matrices. We disabled Deduplication of UMIs (--soloUMIdedup NoDedup) and set strand information to unstranded (--soloStrand unstranded) to match the SmartSeq protocol. We imported Raw counts into Python (v. 3.12.2) using Scanpy (v. 1.10.3). We performed Quality control filtering according to the criteria described in the original study for E-MTAB-5061 dataset(58), with an additional threshold excluding cells with more than 30% of reads mapping to mitochondrial genes, while we excluded cells expressing less than 3000, more than 11000 genes, more than 30% mitochondrial reads and predicted doublets for GSE214517 dataset(59). We used the top 1000 most variable genes for dimensionality reduction and clustering. We performed harmony integration on the GSE214517 dataset to correct for donor-specific effects, whereas we applied no integration to the E-MTAB-5061, as we did not observe donor-driven clustering. We ranked differentially expressed genes between clusters ranked, and we assigned cell types assigned based on the top 5 differentially expressed genes as well as known markers for pancreatic islet subtypes. We reanalyzed the publicly available single-cell dataset GSE153855(59) for pancreatic islets using the preprocessed data deposited in the Gene Expression Omnibus (GEO). We imported the gene expression counts matrix (GSE153855_Expression_counts_HQ_allsamples.txt.gz) and the corresponding cell annotation file (GSE153855_Cell_annotation.txt.gz) into Scanpy (v. 1.10.3). We performed dimensionality reduction and clustering using the top 1000 most variable genes. We used the annotated cell-type labels to verify the expression of established endocrine and exocrine islet marker genes prior to downstream analyses of gene expression across cell types.

### GWAS analysis

Harmonized GWAS summary statistics were downloaded from the GWAS Catalog. Reported credible sets on chromosome 1 were obtained from the Open Targets Genetics platform(64,108,109) and regional association plots were generated using R (v 4.3.0). The most likely causal gene at each locus was prioritised based on the L2G score developed by the Open Targets.

### FLAM-seq libraries

We generated FLAM-seq libraries using the previously established protocol (52), with modifications mentioned below. We used DNase treated total RNA from INS-1E cells and from pancreas (Amsbio cat. #R1234188-50; Thermo Fisher Scientific cat. #QS0621; Takara Bio cat. #636577). For human pancreas samples, we prepared FLAM-seq libraries starting from 3 µg RNA. For INS-1E samples treated with control siRNA or *Tent5a/c* siRNAs and ones subjected to glucose stimulation, we pooled technical triplicates in equimolar ratios for each experimental biological replicate and prepared FLAM-seq libraries starting from 5 µg RNA. We brought RNA samples to a volume of 100 µL and we carried out poly(A) selection using the Dynabeads mRNA direct micro purification kit (Invitrogen cat. #61021) according to the manufacturer’s recommendation. Polyadenylated RNA was GI-tailed using reagents of the USB poly(A) Length Assay Kit (Thermo Fisher Scientific cat. #764551KT) and purified with 1.8X RNAClean XP beads (Beckmann Coulter cat. #A63987). Subsequently we reverse transcribed GI-tailed RNA with the SMARTScribe Reverse Transcriptase kit (Takara Bio cat. #639537) isoTSO and RT primer 1 (listed in XXX). We purified the resulting cDNA with 0.6x AMPure XP beads (Beckmann Coulter cat. #A63881) and PCR-amplified with the Advantage 2 DNA polymerase mix (Clontech cat. #639201). We purified again the libraries with 0.6x AMPure XP beads and submitted them for adapter ligation and PacBio Revio sequencing (Macrogen).

### Overview of the SnakeFLAME analysis pipeline

To enable scalable and efficient analysis of FLAM-seq libraries, we developed a custom, modular analysis pipeline SnakeFLAME, implemented in Snakemake (v8.27.1) using Python (v3.12.8). This pipeline was designed as an updated alternative to the previously published FLAMAnalysis workflow (52) (https://github.com/rajewsky-lab/FLAMAnalysis), which identifies poly(A) tails directly from full-length reads prior to alignment. Our revised strategy modifies both preprocessing and poly(A) detection logic to allow its usage in deeply sequenced libraries/ high read coverage.

### Read preprocessing and filtering

Raw FASTQ files were processed using Cutadapt (via the Snakemake wrapper, v5.9) to identify the expected FLAM-seq characteristic sequence consisting of [A]×10 followed by [G]×9, or its reverse-complementary sequence in independent runs. Cutadapt was configured to detect the expected FLAM-seq motif with a minimum overlap of 19 nt and allowing up to 10% mismatches (-e 0.1, −O 19). Reads lacking this sequence or containing both forward and reverse motifs, likely representing artifacts, were discarded using a custom bash script based on fastq read name identification.

### Read alignment

Filtered reads were aligned to the H.Sapiens reference genome-GRCh38 (hg38) version 47- or the R.Norvegicus reference genome-GRCr8 release 114-using minimap2 (v2.28) with options -ax splice:hq -uf optimized for Pacbio samples. Only uniquely aligned reads were retained for downstream analyses (option --secondary=no).

### Extraction of soft-clipped sequences and poly(A) tail identification

Poly(A) tail detection was restricted to unaligned terminal read segments. Soft-clipped sequences were extracted from aligned reads by parsing CIGAR strings using custom R scripts executed in a dedicated R environment (v4.4.2) with Bioconductor (Biostrings v2.74.0, GenomicAlignments, GenomicRanges) and tidyverse-compatible packages (dplyr, stringr).

Soft clipped sequences were converted to a DNAString object to facilitate efficient searching and handling of nucleotide sequences. The 5’ and 3’ soft-clipped sequences were concatenated with a delimiter (‘-’) to allow simultaneous analysis. Candidate poly(A) tails were initially identified using pattern matching with vmatchPattern, searching for [A]×7 followed by [G]×9, allowing 2 mismatches, consistent with the expected library design. Reads matching the reverse-complement motif ([C]×9 followed by [T]×7) were reverse-complemented to unify orientation. Entries containing multiple poly(A) and/or poly(T) patterns, or lacking any detectable motif, were excluded. Downstream poly(G) stretches (≥8 consecutive Gs, allowing one mismatch) were used to delimit the start of poly(A) tails, and reads lacking exactly one polyG stretch were discarded. vmatchPattern was further used to identify capped sequences by presence of extra [G] in adaptor sequence.

Poly(A) stretches were further refined by detecting short adenine seed motifs (≥3 consecutive As) within the soft-clipped sequences and iteratively extending them to define the maximal contiguous poly(A) tail. This approach increases speed, as the extension is performed by seed motifs rather than by individual nucleotides.

Final poly(A) calls were retained only if the tail exceeded a minimum length (>7 nt) and consisted predominantly of adenines (A-frequency ≥0.85). The final poly(A) stretch was restricted to contiguous adenines and did not include the concatenation delimiter (‘–’) separating 5’ and 3’ soft-clipped segments. Stretches with lower adenine frequency were excluded as likely representing merged cDNA molecules or artifacts.

### Isoform quantification and strand validation

Reads with confirmed poly(A) status were selected by read name using samtools view (v1.21). Transcript-level isoform quantification and annotation were performed using IsoQuant (v3.6.2) using *H.Sapiens* reference genome-GRCh38 (hg38) version 47- or the *R.Norvegicus* reference genome-GRCr8 release 114.

Poly(A) calls were integrated with IsoQuant transcript assignments information to validate strand consistency. For each read, the inferred poly(A) orientation was compared with the annotated transcript strand, and reads were classified as PASS or FAIL accordingly. Lists of read name identifiers corresponding to each category were generated for downstream filtering.

### Filtering of aligned reads and downstream quantification

Aligned BAM files were further filtered to retain only reads with “PASS” poly(A) calls, followed by sorting and indexing. Transcript- and gene-level quantification were repeated on the filtered BAM files, enabling analyses restricted to reads with confidently assigned poly(A) tails.

### Reproducibility and software availability

The workflow is fully reproducible, version-controlled, and supports parallel execution on a high-performance computing cluster. All scripts, environments, and workflows are available at https://github.com/Legnini-Group/SnakeFLAME/.

### Visualization of FLAM-seq alignments

FLAM-seq alignments including their poly(A) tails were visualized in genome browsers (UCSC), taking advantage of the fact that mismatched bases are annotated with a different colour. To achieve this, first soft clipped sequences which contain both poly(A) tails and adapters were trimmed from the read alignments, poly(A) tail sequences were extracted from the output of the snakemake pipeline (polyA_stretch_clipped column in *polyA_final_pass.txt file) and subsequently appended to the trimmed reads, modifying the CIGAR string accordingly.

## Footnotes

1 From here on, lower case notation is used to indicate rodent genes studied in rat INS1-E cells, as compared to human upper case notation.

## References

1. Giddings SJ, Chirgwin J, Permutt MA. Evaluation of rat insulin messenger RNA in pancreatic and extrapancreatic tissues. Diabetologia. 1985 June;28(6):343–7.

2. Brunstedt J, Chan SJ. Direct effect of glucose on the preproinsulin mRNA level in isolated pancreatic islets. Biochem Biophys Res Commun. 1982 June 30;106(4):1383–9.

3. Hammonds P, Schofield PN, Ashcroft SJ, Sutton R, Gray DW. Regulation and specificity of glucose-stimulated insulin gene expression in human islets of Langerhans. FEBS Lett. 1987 Oct 19;223(1):131–7.

4. Eizirik DL, Korbutt GS, Hellerström C. Prolonged exposure of human pancreatic islets to high glucose concentrations in vitro impairs the beta-cell function. J Clin Invest. 1992 Oct;90(4):1263–8.

5. Ling Z, Kiekens R, Mahler T, Schuit FC, Pipeleers-Marichal M, Sener A, et al. Effects of chronically elevated glucose levels on the functional properties of rat pancreatic beta-cells. Diabetes. 1996 Dec;45(12):1774–82.

6. Schuit FC, Kiekens R, Pipeleers DG. Measuring the balance between insulin synthesis and insulin release. Biochem Biophys Res Commun. 1991 Aug 15;178(3):1182–7.

7. Ehehalt F, Knoch K, Erdmann K, Krautz C, Jäger M, Steffen A, et al. Impaired insulin turnover in islets from type 2 diabetic patients. Islets. 2010 Jan;2(1):30–6.

8. Nielsen DA, Welsh M, Casadaban MJ, Steiner DF. Control of insulin gene expression in pancreatic beta-cells and in an insulin-producing cell line, RIN-5F cells. I. Effects of glucose and cyclic AMP on the transcription of insulin mRNA. J Biol Chem. 1985 Nov 5;260(25):13585–9.

9. Leibiger B, Moede T, Schwarz T, Brown GR, Köhler M, Leibiger IB, et al. Short-term regulation of insulin gene transcription by glucose. Proc Natl Acad Sci U S A. 1998 Aug 4;95(16):9307–12.

10. Furukawa N, Shirotani T, Nakamaru K, Matsumoto K, Shichiri M, Araki E. Regulation of the insulin gene transcription by glucose. Endocr J. 2002 Apr;49(2):121–30.

11. Kataoka K, Han S-I, Shioda S, Hirai M, Nishizawa M, Handa H. MafA is a glucose-regulated and pancreatic beta-cell-specific transcriptional activator for the insulin gene. J Biol Chem. 2002 Dec 20;277(51):49903–10.

12. Park K-G, Lee K-M, Seo H-Y, Suh J-H, Kim H-S, Wang L, et al. Glucotoxicity in the INS-1 rat insulinoma cell line is mediated by the orphan nuclear receptor small heterodimer partner. Diabetes. 2007 Feb;56(2):431–7.

13. Olson LK, Redmon JB, Towle HC, Robertson RP. Chronic exposure of HIT cells to high glucose concentrations paradoxically decreases insulin gene transcription and alters binding of insulin gene regulatory protein. J Clin Invest. 1993 July;92(1):514–9.

14. Olson LK, Sharma A, Peshavaria M, Wright CV, Towle HC, Rodertson RP, et al. Reduction of insulin gene transcription in HIT-T15 beta cells chronically exposed to a supraphysiologic glucose concentration is associated with loss of STF-1 transcription factor expression. Proc Natl Acad Sci U S A. 1995 Sept 26;92(20):9127–31.

15. Itoh N, Okamoto H. Translational control of proinsulin synthesis by glucose. Nature. 1980 Jan 3;283(5742):100–2.

16. Wicksteed B, Uchizono Y, Alarcon C, McCuaig JF, Shalev A, Rhodes CJ. A cis-element in the 5’ untranslated region of the preproinsulin mRNA (ppIGE) is required for glucose regulation of proinsulin translation. Cell Metab. 2007 Mar;5(3):221–7.

17. Wicksteed B, Herbert TP, Alarcon C, Lingohr MK, Moss LG, Rhodes CJ. Cooperativity between the preproinsulin mRNA untranslated regions is necessary for glucose-stimulated translation. J Biol Chem. 2001 June 22;276(25):22553–8.

18. Knoch K-P, Bergert H, Borgonovo B, Saeger H-D, Altkrüger A, Verkade P, et al. Polypyrimidine tract-binding protein promotes insulin secretory granule biogenesis. Nat Cell Biol. 2004 Mar;6(3):207–14.

19. Muralidharan B, Bakthavachalu B, Pathak A, Seshadri V. A minimal element in 5’UTR of insulin mRNA mediates its translational regulation by glucose. FEBS Lett. 2007 Aug 21;581(21):4103–8.

20. Lee EK, Kim W, Tominaga K, Martindale JL, Yang X, Subaran SS, et al. RNA-binding protein HuD controls insulin translation. Mol Cell. 2020 Dec 17;80(6):1135.

21. Fred RG, Mehrabi S, Adams CM, Welsh N. PTB and TIAR binding to insulin mRNA 3’-and 5’UTRs; implications for insulin biosynthesis and messenger stability. Heliyon. 2016 Sept;2(9):e00159.

22. Cheruiyot A, Hollister-Lock J, Sullivan B, Pan H, Dreyfuss JM, Bonner-Weir S, et al. Sustained hyperglycemia specifically targets translation of mRNAs for insulin secretion. J Clin Invest [Internet]. 2023 Nov 30;134(3). Available from: 10.1172/JCI173280

23. Hassler JR, Scheuner DL, Wang S, Han J, Kodali VK, Li P, et al. The IRE1α/XBP1s pathway is essential for the glucose response and protection of β cells. PLoS Biol. 2015 Oct;13(10):e1002277.

24. Scheuner D, Vander Mierde D, Song B, Flamez D, Creemers JWM, Tsukamoto K, et al. Control of mRNA translation preserves endoplasmic reticulum function in beta cells and maintains glucose homeostasis. Nat Med. 2005 July;11(7):757–64.

25. Lipson KL, Fonseca SG, Ishigaki S, Nguyen LX, Foss E, Bortell R, et al. Regulation of insulin biosynthesis in pancreatic beta cells by an endoplasmic reticulum-resident protein kinase IRE1. Cell Metab. 2006 Sept;4(3):245–54.

26. Welsh M, Nielsen DA, MacKrell AJ, Steiner DF. Control of insulin gene expression in pancreatic beta-cells and in an insulin-producing cell line, RIN-5F cells. II. Regulation of insulin mRNA stability. J Biol Chem. 1985 Nov 5;260(25):13590–4.

27. Tillmar L, Carlsson C, Welsh N. Control of insulin mRNA stability in rat pancreatic islets. Regulatory role of a 3’-untranslated region pyrimidine-rich sequence. J Biol Chem. 2002 Jan 11;277(2):1099–106.

28. Kühn U, Gündel M, Knoth A, Kerwitz Y, Rüdel S, Wahle E. Poly(A) tail length is controlled by the nuclear poly(A)-binding protein regulating the interaction between poly(A) polymerase and the cleavage and polyadenylation specificity factor. J Biol Chem. 2009 Aug 21;284(34):22803–14.

29. Keller RW, Kühn U, Aragón M, Bornikova L, Wahle E, Bear DG. The nuclear poly(A) binding protein, PABP2, forms an oligomeric particle covering the length of the poly(A) tail. J Mol Biol. 2000 Mar 31;297(3):569–83.

30. Passmore LA, Coller J. Roles of mRNA poly(A) tails in regulation of eukaryotic gene expression. Nat Rev Mol Cell Biol. 2022 Feb;23(2):93–106.

31. Brown CE, Sachs AB. Poly(A) tail length control in Saccharomyces cerevisiae occurs by message-specific deadenylation. Mol Cell Biol. 1998 Nov;18(11):6548–59.

32. Eisen TJ, Eichhorn SW, Subtelny AO, Lin KS, McGeary SE, Gupta S, et al. The dynamics of cytoplasmic mRNA metabolism. Mol Cell. 2020 Feb 20;77(4):786–799.e10.

33. Czarnocka-Cieciura A, Poznański J, Turtola M, Tomecki R, Krawczyk PS, Mroczek S, et al. Modeling of mRNA deadenylation rates reveal a complex relationship between mRNA deadenylation and decay. EMBO J. 2024 Dec;43(24):6525–54.

34. Yamashita A, Chang T-C, Yamashita Y, Zhu W, Zhong Z, Chen C-YA, et al. Concerted action of poly(A) nucleases and decapping enzyme in mammalian mRNA turnover. Nat Struct Mol Biol. 2005 Dec;12(12):1054–63.

35. Chen C-YA, Shyu A-B. Mechanisms of deadenylation-dependent decay. Wiley Interdiscip Rev RNA. 2011 Mar;2(2):167–83.

36. Kuchta K, Knizewski L, Wyrwicz LS, Rychlewski L, Ginalski K. Comprehensive classification of nucleotidyltransferase fold proteins: identification of novel families and their representatives in human. Nucleic Acids Res. 2009 Dec;37(22):7701–14.

37. Kuchta K, Muszewska A, Knizewski L, Steczkiewicz K, Wyrwicz LS, Pawlowski K, et al. FAM46 proteins are novel eukaryotic non-canonical poly(A) polymerases. Nucleic Acids Res. 2016 May 5;44(8):3534–48.

38. Lacidogna D, Pennacchio S, Milan E. TENT5/FAM46: An enigmatic family of secretory tuners. Traffic. 2025 Apr;26(4–6):e70011.

39. Bilska A, Kusio-Kobiałka M, Krawczyk PS, Gewartowska O, Tarkowski B, Kobyłecki K, et al. Immunoglobulin expression and the humoral immune response is regulated by the non-canonical poly(A) polymerase TENT5C. Nat Commun. 2020 Apr 27;11(1):2032.

40. Krawczyk PS, Mazur M, Orzeł W, Gewartowska O, Jeleń S, Antczak W, et al. Re-adenylation by TENT5A enhances efficacy of SARS-CoV-2 mRNA vaccines. Nature. 2025 May;641(8064):984–92.

41. Weyer C, Bogardus C, Mott DM, Pratley RE. The natural history of insulin secretory dysfunction and insulin resistance in the pathogenesis of type 2 diabetes mellitus. J Clin Invest. 1999 Sept;104(6):787–94.

42. Arunagiri A, Haataja L, Pottekat A, Pamenan F, Kim S, Zeltser LM, et al. Proinsulin misfolding is an early event in the progression to type 2 diabetes. Elife [Internet]. 2019 June 11;8. Available from: 10.7554/eLife.44532

43. Yamane S, Hamamoto Y, Harashima S-I, Harada N, Hamasaki A, Toyoda K, et al. GLP-1 receptor agonist attenuates endoplasmic reticulum stress-mediated β-cell damage in Akita mice. J Diabetes Investig. 2011 Apr 7;2(2):104–10.

44. Yusta B, Baggio LL, Estall JL, Koehler JA, Holland DP, Li H, et al. GLP-1 receptor activation improves beta cell function and survival following induction of endoplasmic reticulum stress. Cell Metab. 2006 Nov;4(5):391–406.

45. Yong J, Parekh VS, Reilly SM, Nayak J, Chen Z, Lebeaupin C, et al. Chop/Ddit3 depletion in β cells alleviates ER stress and corrects hepatic steatosis in mice. Sci Transl Med. 2021 July 28;13(604):eaba9796.

46. Szabat M, Page MM, Panzhinskiy E, Skovsø S, Mojibian M, Fernandez-Tajes J, et al. Reduced insulin production relieves endoplasmic reticulum stress and induces β cell proliferation. Cell Metab. 2016 Jan 12;23(1):179–93.

47. Templeman NM, Clee SM, Johnson JD. Suppression of hyperinsulinaemia in growing female mice provides long-term protection against obesity. Diabetologia. 2015 Oct;58(10):2392–402.

48. Page MM, Skovsø S, Cen H, Chiu AP, Dionne DA, Hutchinson DF, et al. Reducing insulin via conditional partial gene ablation in adults reverses diet-induced weight gain. FASEB J. 2018 Mar;32(3):1196–206.

49. Fucci C, Resnati M, Riva E, Perini T, Ruggieri E, Orfanelli U, et al. The interaction of the tumor suppressor FAM46C with p62 and FNDC3 proteins integrates protein and secretory homeostasis. Cell Rep. 2020 Sept 22;32(12):108162.

50. Mroczek S, Chlebowska J, Kuliński TM, Gewartowska O, Gruchota J, Cysewski D, et al. The non-canonical poly(A) polymerase FAM46C acts as an onco-suppressor in multiple myeloma. Nat Commun. 2017 Sept 20;8(1):619.

51. Zhu YX, Shi C-X, Bruins LA, Jedlowski P, Wang X, Kortüm KM, et al. Loss of FAM46C promotes cell survival in myeloma. Cancer Res. 2017 Aug 15;77(16):4317–27.

52. Legnini I, Alles J, Karaiskos N, Ayoub S, Rajewsky N. FLAM-seq: full-length mRNA sequencing reveals principles of poly(A) tail length control. Nat Methods. 2019 Sept;16(9):879–86.

53. Brouze M, Czarnocka-Cieciura A, Gewartowska O, Kusio-Kobiałka M, Jachacy K, Szpila M, et al. TENT5-mediated polyadenylation of mRNAs encoding secreted proteins is essential for gametogenesis in mice. Nat Commun. 2024 June 22;15(1):5331.

54. Alles J, Legnini I, Pacelli M, Rajewsky N. Rapid nuclear deadenylation of mammalian messenger RNA. iScience. 2023 Jan 20;26(1):105878.

55. Lizio M, Abugessaisa I, Noguchi S, Kondo A, Hasegawa A, Hon CC, et al. Update of the FANTOM web resource: expansion to provide additional transcriptome atlases. Nucleic Acids Res. 2019 Jan 8;47(D1):D752–8.

56. Lizio M, Harshbarger J, Shimoji H, Severin J, Kasukawa T, Sahin S, et al. Gateways to the FANTOM5 promoter level mammalian expression atlas. Genome Biol. 2015 Jan 5;16(1):22.

57. GTEx Consortium. The GTEx Consortium atlas of genetic regulatory effects across human tissues. Science. 2020 Sept 11;369(6509):1318–30.

58. Segerstolpe Å, Palasantza A, Eliasson P, Andersson E-M, Andréasson A-C, Sun X, et al. Single-cell transcriptome profiling of human pancreatic islets in health and type 2 diabetes. Cell Metab. 2016 Oct 11;24(4):593–607.

59. Martínez-López JA, Lindqvist A, Lopez-Pascual A, Harder A, Chen P, Ngara M, et al. Single-cell mRNA-regulation analysis reveals cell type-specific mechanisms of type 2 diabetes. Nat Commun. 2025 Oct 27;16(1):9475.

60. Ziegenhain C, Vieth B, Parekh S, Reinius B, Guillaumet-Adkins A, Smets M, et al. Comparative analysis of single-cell RNA sequencing methods. Mol Cell. 2017 Feb 16;65(4):631–643.e4.

61. Soneson C, Robinson MD. Bias, robustness and scalability in single-cell differential expression analysis. Nat Methods. 2018 Apr;15(4):255–61.

62. Kaestner KH, Powers AC, Naji A, HPAP Consortium, Atkinson MA. NIH initiative to improve understanding of the pancreas, islet, and autoimmunity in type 1 diabetes: The Human Pancreas Analysis Program (HPAP). Diabetes. 2019 July;68(7):1394–402.

63. Shapira SN, Naji A, Atkinson MA, Powers AC, Kaestner KH. Understanding islet dysfunction in type 2 diabetes through multidimensional pancreatic phenotyping: The Human Pancreas Analysis Program. Cell Metab. 2022 Dec 6;34(12):1906–13.

64. Buniello A, Suveges D, Cruz-Castillo C, Llinares MB, Cornu H, Lopez I, et al. Open Targets Platform: facilitating therapeutic hypotheses building in drug discovery. Nucleic Acids Res. 2025 Jan 6;53(D1):D1467–75.

65. Verma A, Huffman JE, Rodriguez A, Conery M, Liu M, Ho Y-L, et al. Diversity and scale: Genetic architecture of 2068 traits in the VA Million Veteran Program. Science. 2024 July 19;385(6706):eadj1182.

66. Butler AE, Janson J, Bonner-Weir S, Ritzel R, Rizza RA, Butler PC. Beta-cell deficit and increased beta-cell apoptosis in humans with type 2 diabetes. Diabetes. 2003 Jan;52(1):102–10.

67. Silva PN, Mayrhofer JE, Potalitsyn P, Trüllinger GO, Godbersen S, Amstutz N, et al. Polyadenylation of insulin mRNA by Tent5a regulates pancreatic beta cells. Nat Commun [Internet]. 2026 May 20; Available from: 10.1038/s41467-026-72905-8

68. Knappskog S, Ravneberg H, Gjerdrum C, Trösse C, Stern B, Pryme IF. The level of synthesis and secretion of Gaussia princeps luciferase in transfected CHO cells is heavily dependent on the choice of signal peptide. J Biotechnol. 2007 Mar 10;128(4):705–15.

69. Liu J, O’Kane DJ, Escher A. Secretion of functional Renilla reniformis luciferase by mammalian cells. Gene. 1997 Dec 12;203(2):141–8.

70. Schmidt SF, Madsen JGS, Frafjord KØ, Poulsen L la C, Salö S, Boergesen M, et al. Integrative genomics outlines a biphasic glucose response and a ChREBP-RORγ axis regulating proliferation in β cells. Cell Rep. 2016 Aug 30;16(9):2359–72.

71. Quezada E, Knoch K-P, Vasiljevic J, Seiler A, Pal A, Gunasekaran A, et al. Aldolase-regulated G3BP1/2+ condensates control insulin mRNA storage in beta cells. EMBO J. 2025 July;44(13):3669–96.

72. Back SH, Scheuner D, Han J, Song B, Ribick M, Wang J, et al. Translation attenuation through eIF2alpha phosphorylation prevents oxidative stress and maintains the differentiated state in beta cells. Cell Metab. 2009 July;10(1):13–26.

73. Scheuner D, Song B, McEwen E, Liu C, Laybutt R, Gillespie P, et al. Translational control is required for the unfolded protein response and in vivo glucose homeostasis. Mol Cell. 2001 June;7(6):1165–76.

74. Pirot P, Naamane N, Libert F, Magnusson NE, Ørntoft TF, Cardozo AK, et al. Global profiling of genes modified by endoplasmic reticulum stress in pancreatic beta cells reveals the early degradation of insulin mRNAs. Diabetologia. 2007 May;50(5):1006–14.

75. Lipson KL, Ghosh R, Urano F. The role of IRE1alpha in the degradation of insulin mRNA in pancreatic beta-cells. PLoS One. 2008 Feb 20;3(2):e1648.

76. van Tienhoven R, O’Meally D, Scott TA, Morris KV, Williams JC, Kaddis JS, et al. Genetic protection from type 1 diabetes resulting from accelerated insulin mRNA decay. Cell. 2025 May 1;188(9):2407–2416.e9.

77. Gray NK, Coller JM, Dickson KS, Wickens M. Multiple portions of poly(A)-binding protein stimulate translation in vivo. EMBO J. 2000 Sept 1;19(17):4723–33.

78. Tucker M, Valencia-Sanchez MA, Staples RR, Chen J, Denis CL, Parker R. The transcription factor associated Ccr4 and Caf1 proteins are components of the major cytoplasmic mRNA deadenylase in Saccharomyces cerevisiae. Cell. 2001 Feb 9;104(3):377–86.

79. Goldstrohm AC, Wickens M. Multifunctional deadenylase complexes diversify mRNA control. Nat Rev Mol Cell Biol. 2008 Apr;9(4):337–44.

80. Eichhorn SW, Subtelny AO, Kronja I, Kwasnieski JC, Orr-Weaver TL, Bartel DP. mRNA poly(A)-tail changes specified by deadenylation broadly reshape translation in Drosophila oocytes and early embryos. Elife [Internet]. 2016 July 30;5. Available from: 10.7554/eLife.16955

81. Subtelny AO, Eichhorn SW, Chen GR, Sive H, Bartel DP. Poly(A)-tail profiling reveals an embryonic switch in translational control. Nature. 2014 Apr 3;508(7494):66–71.

82. Muhlrad D, Decker CJ, Parker R. Deadenylation of the unstable mRNA encoded by the yeast MFA2 gene leads to decapping followed by 5’-->3’ digestion of the transcript. Genes Dev. 1994 Apr 1;8(7):855–66.

83. Baptissart M, Gupta A, Poirot AC, Papas BN, Morgan M. TENT5C extends Odf1 poly(A) tail to sustain sperm morphogenesis and fertility [Internet]. bioRxivorg. 2025. Available from: 10.1101/2025.03.20.644152

84. Gewartowska O, Aranaz-Novaliches G, Krawczyk PS, Mroczek S, Kusio-Kobiałka M, Tarkowski B, et al. Cytoplasmic polyadenylation by TENT5A is required for proper bone formation. Cell Rep. 2021 Apr 20;35(3):109015.

85. Mazur M, Gumińska N, Brouze A, Cysewski D, Mleczko-Sanecka K, Niklewicz M, et al. Efficient globin production during terminal erythropoiesis depends on the synergistic action of TENT5C poly(A) polymerase and LARP4/5 [Internet]. bioRxiv. bioRxiv; 2024. Available from: 10.1101/2024.11.14.623596

86. Madsen AL, Bonàs-Guarch S, Gheibi S, Prasad R, Vangipurapu J, Ahuja V, et al. Genetic architecture of oral glucose-stimulated insulin release provides biological insights into type 2 diabetes aetiology. Nat Metab. 2024 Oct;6(10):1897–912.

87. Rottner AK, Ye Y, Navarro-Guerrero E, Rajesh V, Pollner A, Bevacqua RJ, et al. A genome-wide CRISPR screen identifies CALCOCO2 as a regulator of beta cell function influencing type 2 diabetes risk. Nat Genet. 2023 Jan;55(1):54–65.

88. Harding HP, Zeng H, Zhang Y, Jungries R, Chung P, Plesken H, et al. Diabetes mellitus and exocrine pancreatic dysfunction in perk-/- mice reveals a role for translational control in secretory cell survival. Mol Cell. 2001 June;7(6):1153–63.

89. Usui M, Yamaguchi S, Tanji Y, Tominaga R, Ishigaki Y, Fukumoto M, et al. Atf6α-null mice are glucose intolerant due to pancreatic β-cell failure on a high-fat diet but partially resistant to diet-induced insulin resistance. Metabolism. 2012 Aug;61(8):1118–28.

90. Ladiges WC, Knoblaugh SE, Morton JF, Korth MJ, Sopher BL, Baskin CR, et al. Pancreatic beta-cell failure and diabetes in mice with a deletion mutation of the endoplasmic reticulum molecular chaperone gene P58IPK. Diabetes. 2005 Apr;54(4):1074–81.

91. Sharma RB, O’Donnell AC, Stamateris RE, Ha B, McCloskey KM, Reynolds PR, et al. Insulin demand regulates β cell number via the unfolded protein response. J Clin Invest. 2015 Oct 1;125(10):3831–46.

92. Yong J, Johnson JD, Arvan P, Han J, Kaufman RJ. Therapeutic opportunities for pancreatic β-cell ER stress in diabetes mellitus. Nat Rev Endocrinol. 2021 Aug;17(8):455–67.

93. Lytrivi M, Tong Y, Virgilio E, Yi X, Cnop M. Diabetes mellitus and the key role of endoplasmic reticulum stress in pancreatic β cells. Nat Rev Endocrinol. 2025 Sept;21(9):546–63.

94. Song B, Scheuner D, Ron D, Pennathur S, Kaufman RJ. Chop deletion reduces oxidative stress, improves beta cell function, and promotes cell survival in multiple mouse models of diabetes. J Clin Invest. 2008 Oct;118(10):3378–89.

95. Groza T, Gomez FL, Mashhadi HH, Muñoz-Fuentes V, Gunes O, Wilson R, et al. The International Mouse Phenotyping Consortium: comprehensive knockout phenotyping underpinning the study of human disease. Nucleic Acids Res. 2023 Jan 6;51(D1):D1038–45.

96. Steinbrecht D, Minia I, Milek M, Meisig J, Blüthgen N, Landthaler M. Subcellular mRNA kinetic modeling reveals nuclear retention as rate-limiting. Mol Syst Biol. 2024 Dec;20(12):1346–71.

97. Laemmli UK. Cleavage of structural proteins during the assembly of the head of bacteriophage T4. Nature. 1970 Aug 15;227(5259):680–5.

98. Müller T, Kalxdorf M, Longuespée R, Kazdal DN, Stenzinger A, Krijgsveld J. Automated sample preparation with SP3 for low-input clinical proteomics. Mol Syst Biol. 2020 Jan;16(1):e9111.

99. Guzman UH, Martinez-Val A, Ye Z, Damoc E, Arrey TN, Pashkova A, et al. Ultra-fast label-free quantification and comprehensive proteome coverage with narrow-window data-independent acquisition. Nat Biotechnol. 2024 Dec;42(12):1855–66.

100. Demichev V, Messner CB, Vernardis SI, Lilley KS, Ralser M. DIA-NN: neural networks and interference correction enable deep proteome coverage in high throughput. Nat Methods. 2020 Jan;17(1):41–4.

101. Välikangas T, Suomi T, Elo LL. A systematic evaluation of normalization methods in quantitative label-free proteomics. Brief Bioinform. 2018 Jan 1;19(1):1–11.

102. Stekhoven DJ, Bühlmann P. MissForest--non-parametric missing value imputation for mixed-type data. Bioinformatics. 2012 Jan 1;28(1):112–8.

103. Harris L, Fondrie WE, Oh S, Noble WS. Evaluating proteomics imputation methods with improved criteria. J Proteome Res. 2023 Nov 3;22(11):3427–38.

104. Di Tommaso P, Chatzou M, Floden EW, Barja PP, Palumbo E, Notredame C. Nextflow enables reproducible computational workflows. Nat Biotechnol. 2017 Apr 11;35(4):316–9.

105. Love MI, Huber W, Anders S. Moderated estimation of fold change and dispersion for RNA-seq data with DESeq2. Genome Biol. 2014;15(12):550.

106. Dobin A, Davis CA, Schlesinger F, Drenkow J, Zaleski C, Jha S, et al. STAR: ultrafast universal RNA-seq aligner. Bioinformatics. 2013 Jan 1;29(1):15–21.

107. Patro R, Duggal G, Love MI, Irizarry RA, Kingsford C. Salmon provides fast and bias-aware quantification of transcript expression. Nat Methods. 2017 Apr;14(4):417–9.

108. National Center for Biotechnology Information. Index of /dbgap/studies/phs002453/phs002453.v1.p1/analyses/GIA [Internet]. [cited 2026 Feb 25]. Available from: https://ftp.ncbi.nlm.nih.gov/dbgap/studies/phs002453/phs002453.v1.p1/analyses/GIA/

109. European Molecular Biology Laboratory, European Bioinformatics Institute. Index of /pub/databases/gwas/summary_statistics/GCST90475001-GCST90476000/GCST90475667/harmonised [Internet]. [cited 2026 Feb 25]. Available from: http://ftp.ebi.ac.uk/pub/databases/gwas/summary_statistics/GCST90475001-GCST90476000/GCST90475667/harmonised/

110. Teufel F, Almagro Armenteros JJ, Johansen AR, Gíslason MH, Pihl SI, Tsirigos KD, et al. SignalP 6.0 predicts all five types of signal peptides using protein language models. Nat Biotechnol. 2022 July;40(7):1023–5.

111. Wang R, Ren C, Gao T, Li H, Bo X, Zhu D, et al. SEPDB: a database of secreted proteins. Database (Oxford) [Internet]. 2024 Feb 12;2024. Available from: 10.1093/database/baae007

