## Supplementary Information for "Insulin synthesis is sustained by Tent5 poly(A) polymerases"

Content: supplementary figures and legends 1-5, supplementary table 1.

### Supplementary figures and legends.

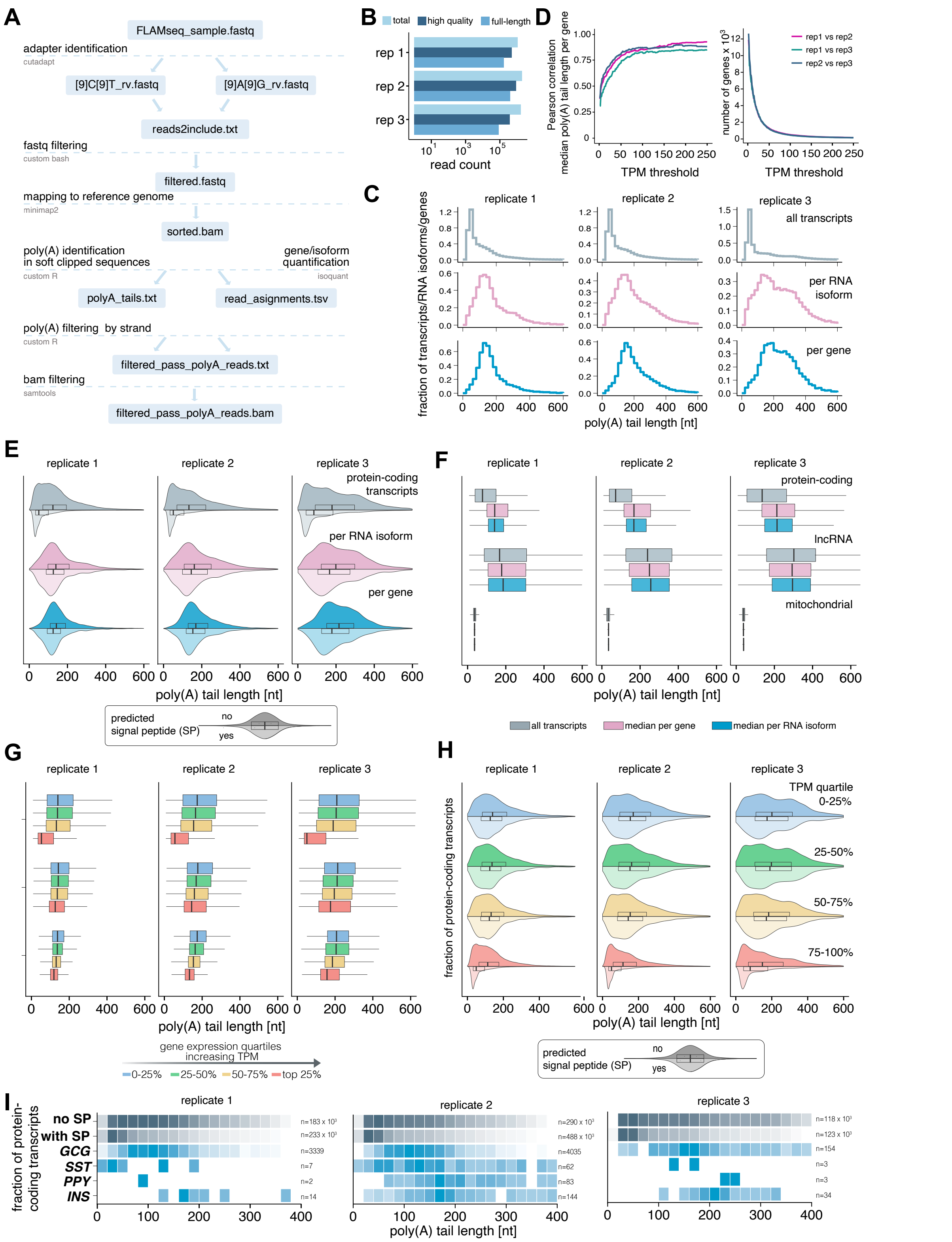


### Supplementary 1. A subset of secretory mRNAs has unusually long poly(A) tails.

A. Schematic outline of the computational pipeline SnakeFLAME, used to analyze FLAM-seq datasets.

B. Read counts across FLAM-seq human pancreas datasets (n=3 independent RNA samples). Total reads indicate all sequencing reads, high-quality reads are those passing SnakeFLAME pipeline filters, containing adaptors and a bona-fide poly(A) tail, while full-length reads span entire transcripts, including the 5'cap as illustrated by the example in Figure 1B.

C. Poly(A) tail length distributions in the single human pancreas datasets (n=3 independent RNA samples) across all detected transcripts (top) or summarized as median poly(A) tail length per gene and per RNA isoform (bottom).

D. Relationship between transcript abundance and reproducibility of gene-level median poly(A) tail length estimates**.** Left, Pearson correlation of gene-level median poly(A) tail lengths between individual human pancreas datasets, calculated for genes exceeding increasing TPM (Transcripts per million) thresholds. Right, the number of genes retained at each TPM threshold.

E. Poly(A) tail length distributions of protein-coding genes in individual human pancreas datasets, stratified by the abscence (top violin) or the presence (bottom violin) of a predicted or validated signal peptide in at least one RNA isoform of the gene, according to SignalP6.0(110). Shown are distributions across all detected transcripts or summarized as median poly(A) tail length per gene and per RNA isoform.

F. Poly(A) tail lengths in individual human pancreas datasets for nuclear encoded protein-coding genes, long non-coding RNA (lncRNA) and mitochondrial genes across all detected transcripts or summarized as median poly(A) tail length per gene and per RNA isoforms.

G. Poly(A) tail lengths for genes stratified into equal expression quartiles based on transcripts per million (TPM) values obtained in individual human pancreas datasets. Poly(A) tail lengths are shown either across all detected transcripts or as median values summarized at the gene or RNA isoform level.

H. Poly(A) tail length distributions for protein-coding transcripts in individual human pancreas datasets, binned into equal expression quartiles based on transcript per million (TPM) values obtained from FLAM-seq and stratified by the absence (upper violin) or presence (lower violin) of a predicted or validated signal peptide in at least one RNA isoform of the corresponding gene, according to SignalP 6.0(110).

I. Poly(A) tail length distributions of the indicated pancreatic hormones compared to all other protein-coding genes within the same gene expression quartile (upper expression bin 75-100% quartile) in individual human pancreas datasets. Protein coding genes are stratified by the absence or the presence of a predicted or validated signal peptide (SP) in at least one RNA isoform of the corresponding gene, according to SignalP 6.0(110). Distributions are displayed as heatmaps showing the relative frequency of reads across poly(A) tail length bins. For each gene (in blue) or gene category (in grey), the color scale was normalized such that the minimum and maximum color intensities correspond to the 5th and 95th percentiles, respectively, of the read density across all tail-length bins.

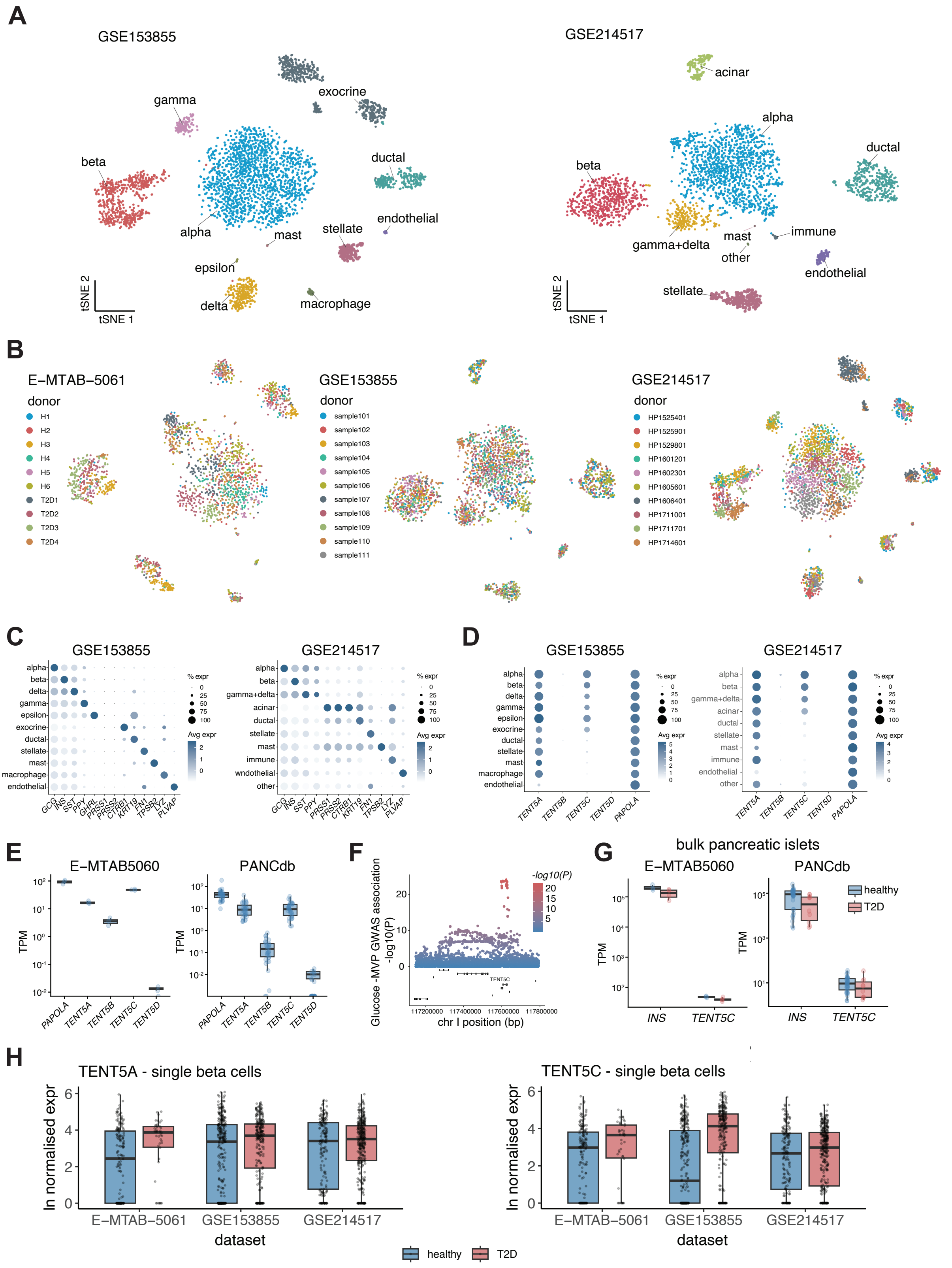


**Supplementary 2. TENT5C is highly expressed in pancreatic beta cells and genetically associated with T2D.**

A. t-SNE projections of GSE153855 (left) and GSE214517 (right) pancreatic islet single-cell RNA-seq datasets, in addition to that shown in the main Fig. 3B. Cells are coloured by annotated cell type identity. Major endocrine populations (Alpha, Beta, Gamma,

Delta), exocrine (Acinar, Ductal), endothelial, and mesenchymal/immune cell populations are resolved in all datasets. For GSE214517, the t-SNE projection was generated from Harmony-integrated principal components to account for donor-specific variation.

B. t-SNE projection of E-MTAB-5061, GSE153855, and GSE214517 datasets coloured by donor/sample identity. Cells from different donors are interspersed across clusters rather than forming donor-specific groupings, indicating minimal donor-driven batch effects. For GSE214517, embeddings were computed from Harmony-integrated principal components to account for donor-specific variation.

C. Dot plot showing the scaled average expression of canonical marker genes across annotated cell populations in GSE153855 (left) and GSE214517 (right). Marker patterns confirm consistent annotation of endocrine, exocrine, mesenchymal/immune, and endothelial populations across datasets.

D. Dot plots showing the average expression of *TENT5A-D* and canonical poly(A) polymerase *PAPOLA* across annotated cell populations in GSE153855 (left) and GSE214517 (right).

E. Expression of *TENT5* family genes in E-MTAB5060 (left) and PANCdb (right) datasets.

F. GWAS association with glucose metabolism at the *TENT5C* locus on chromosome I. Triangle indicates the highest value of association.

G. *INS* and *TENT5C* expression in E-MTAB5060 (left) and PANCdb (right) in healthy and T2D donors.

H. Boxplots showing *TENT5A* and *TENT5C* expression grouped by disease (healthy and T2D) across main pancreatic cell types in E-MTAB-5061, GSE153855, and GSE214517 single-cell RNA-seq datasets. Boxplots show interquartile range and median.

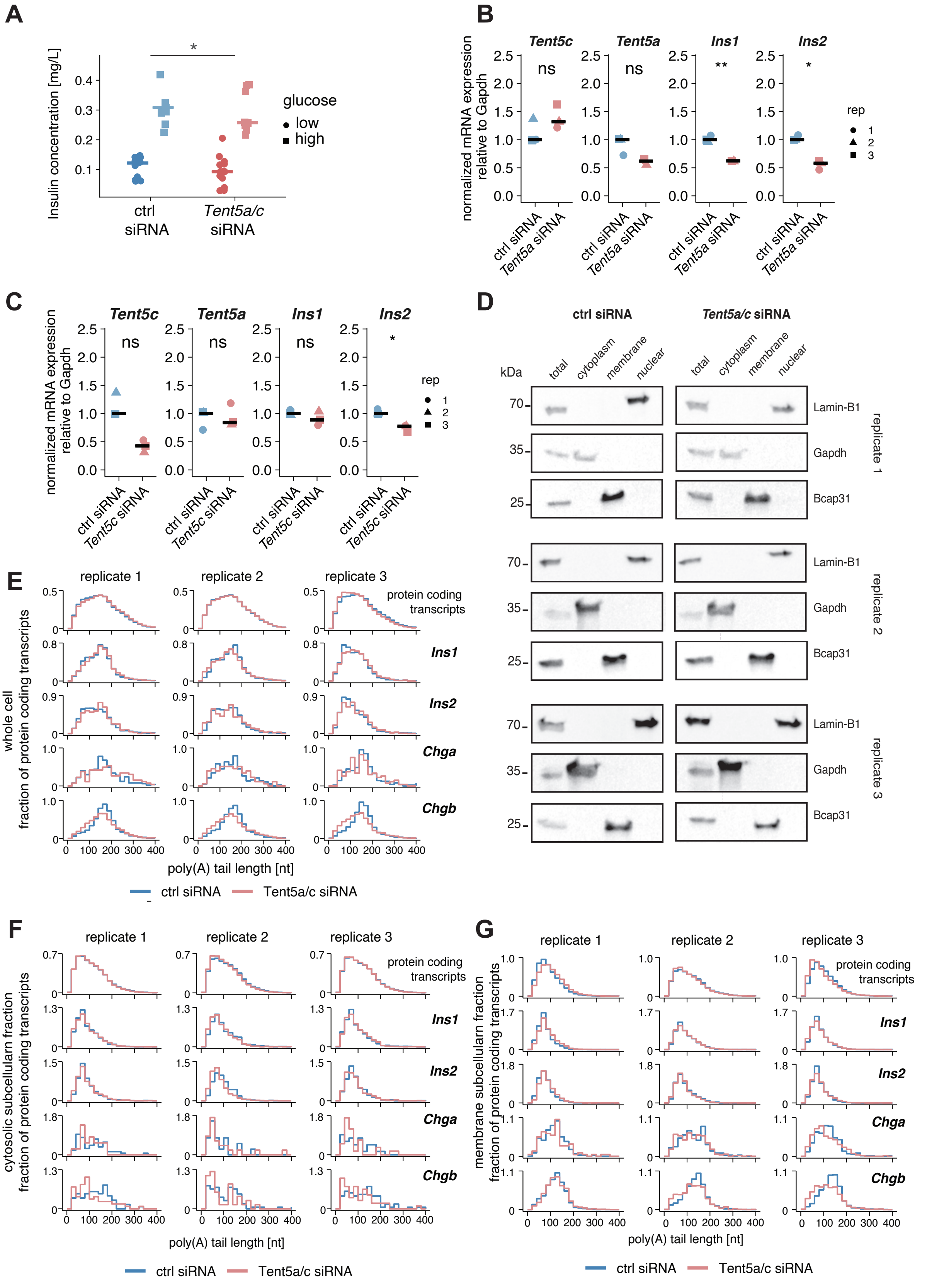


**Supplementary 3. Insulin mRNA abundance is controlled by Tent5a/c poly(A) polymerases.**

A. ELISA measuring insulin secretion upon low glucose (2.8 mM) and high glucose stimulation (28 mM) in INS-1E control cells (ctrl siRNA) and *Tent5a/c* double knockdown cells *(Tent5a/c* siRNA) (n=4 independent experiments, each in n=3 or n=6 technical triplicates). Linear regression was used to test the effect of siRNA treatment on insulin secretion, adjusting for glucose condition and experimental batch (p-value <0,05 (*)).

B. qRT-PCR analysis of the indicated genes in INS-1E control cells (ctrl siRNA) and *Tent5a* knockdown cells *(Tent5a* siRNA). Expression was normalized to *Gapdh* and to control cells. Data represent n = 3 independent experiments, each with technical triplicates; crossbars indicate medians. Paired t-test p-values are indicated as follows: <0.05 (*), <0.01 (**), ns (non-significant).

C. qRT-PCR analysis of the indicated genes in INS-1E control cells (ctrl siRNA) and *Tent5c* knockdown cells *(Tent5c* siRNA). Expression was normalized to *Gapdh* and to control cells. Data represent n = 3 independent experiments, each with technical triplicates; crossbars indicate medians. Paired t-test p-values are indicated as follows: <0.05 (*), ns (non-significant).

D. Western blot analysis to analyze efficiency of subcellular fractionation in INS1-E control cells (ctrl siRNA) or *Tent5a/c* double knockdown (*Tent5a/c* siRNA) for nuclear (Lamin-B1), cytosolic (Gapdh), endoplasmic reticulum (Bcap31) markers. (n=3 independent replicates).

E. Poly(A) tail length distributions of all protein coding transcripts or for the indicated genes for INS-1E control cells (ctrl siRNA) or Tent5a/c double knockdown cells (*Tent5a/c* siRNA) across three individual replicates. Distributions are normalized so that the area under each curve equals 1.

F. Poly(A) tail length distributions of all protein coding transcripts or for the indicated genes in cytosolic subcellular fractions for INS-1E control cells (ctrl siRNA) or Tent5a/c double knockdown cells (*Tent5a/c* siRNA) across three individual replicates. Distributions are normalized so that the area under each curve equals 1.

G. Poly(A) tail length distributions of all protein coding transcripts or for the indicated genes in membrane subcellular fractions for INS-1E control cells (ctrl siRNA) or Tent5a/c double knockdown cells (*Tent5a/c* siRNA) across three individual replicates. Distributions are normalized so that the area under each curve equals 1.

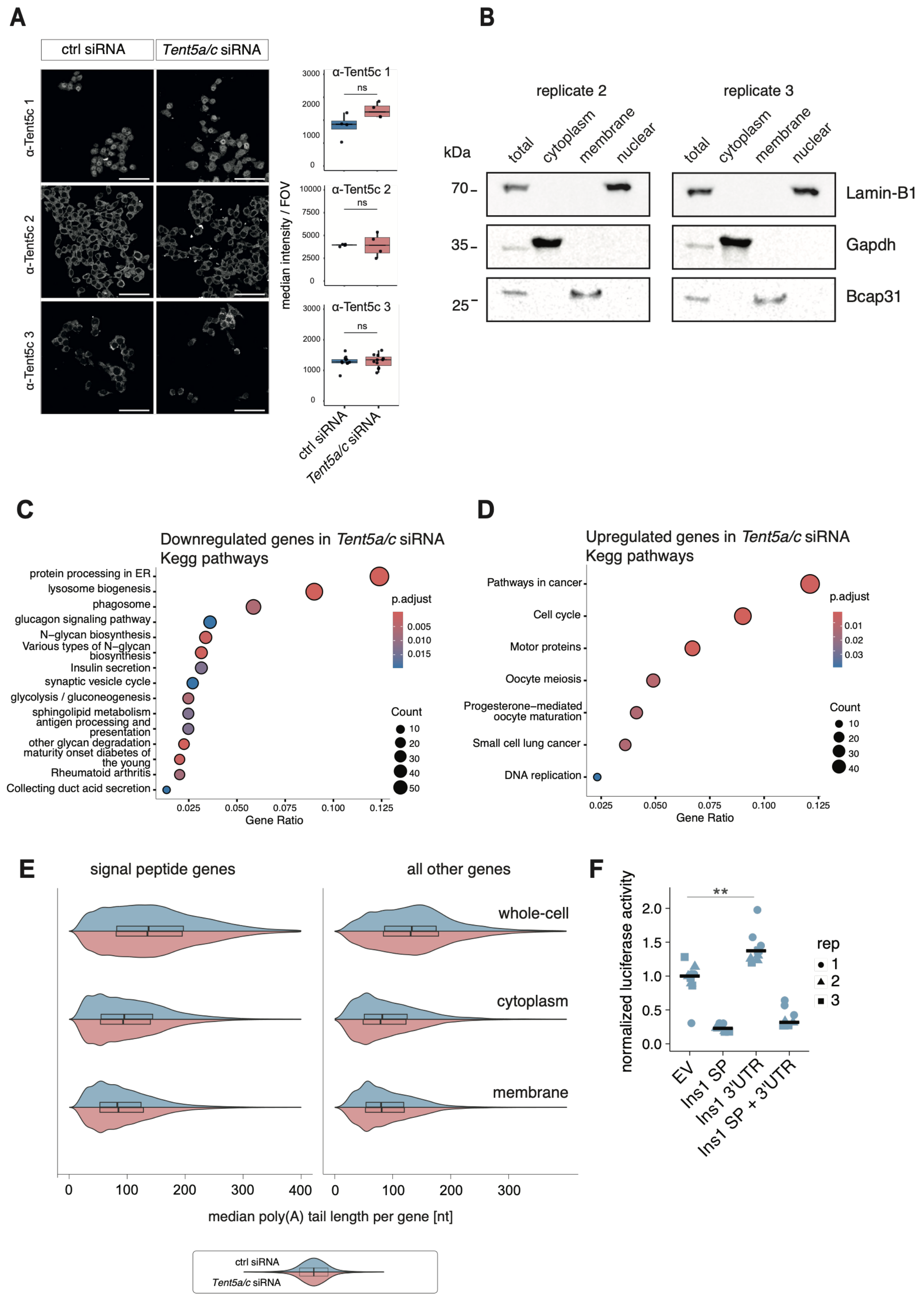


**Supplementary 4. Tent5-mediated regulation of insulin mRNA is stimulated by both its 3'UTR and ER localization.**

A. Immunostaining images for Tent5c staining in INS-1E control cells (ctrl siRNA) or Tent5a/c knockdown cells (*Tent5a/c* siRNA). Three different antibodies recognizing Tent5c were tested (Scale bar = 50 μm). Quantification of the fluorescent signal is shown as median of single-cell mean intensities within each field of view (fov) acquired per condition. Each dot represents one fov. Crossbar indicates medians. Wilcoxon rank-sum test was applied. P-value > 0.05 (ns).

B. Subcellular fractionation of INS-1E cells for subsequent LC-MS proteomics. Western blot analysis measuring nuclear (Lamin-B1), cytosolic (Gapdh), endoplasmic reticulum (Bcap31) markers in INS-1E subcellular fractions and the whole-cell extracts. Two independent replicates in addition to Fig.4C are shown.

C. KEGG pathway enrichment analysis of genes downregulated following *Tent5a/c* knockdown in INS-1E cells. Gene Set Enrichment Analysis (GSEA) was performed on genes significantly downregulated upon Tent5a/c knockdown in bulk RNA-seq data (Fig. 3B). Enriched KEGG pathways are displayed as a dot plot, with dot size indicating the number of genes contributing to each pathway and color representing the adjusted *P* value (FDR).

D. KEGG pathway enrichment analysis of genes upregulated following *Tent5a/c* knockdown in INS-1E cells. Gene Set Enrichment Analysis (GSEA) was performed on genes significantly upregulated upon Tent5a/c knockdown in bulk RNA-seq data (Fig. 3B). Enriched KEGG pathways are displayed as a dot plot, with dot size indicating the number of genes contributing to each pathway and color representing the adjusted *P* value (FDR).

E. Distributions of median poly(A) tail lengths per gene in whole cell, cytoplasmic and membrane fractions from INS1-E control cells (upper violin, ctrl siRNA) *Tent5a/c* knockdown cells (lower violin, *Tent5a/c* siRNA). Genes are stratified by the presence or abscence of a  a signal peptide as predicted for rat genes by SEPDB open (110).

F. Normalized *Renilla luciferase* activities of reporters carrying no fusion (EV), the rat Insulin 1 (*Ins1*) signal peptide (SP), its 3' untranslated region (3'UTR), or both, in control INS-1E cells (ctrl siRNA) (n = 3 independent experiments, each with technical triplicates). *Renilla luciferase* activities were first internally normalized to firefly luciferase activity and then expressed as fold-change relative to EV reporter in control conditions; crossbars indicate medians. Paired t-test p-values are indicated as follows: <0.01 (**).

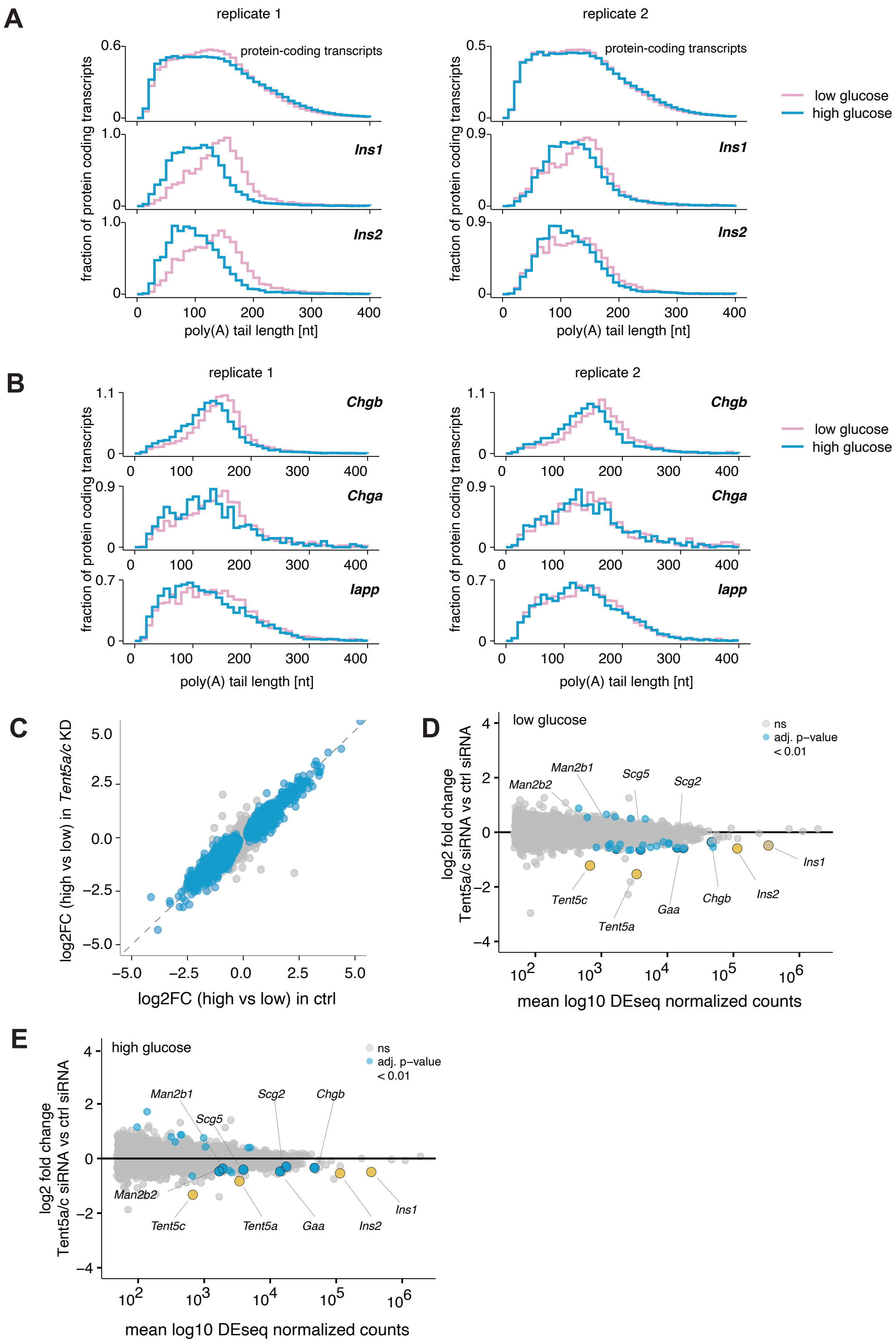


**Supplementary 5. Insulin poly(A) tail shortening and *Tent5a/c* expression are induced by glucose.**

A. Poly(A) tail length distributions for Ins1/2 and all protein coding transcripts upon 1 hour glucose starvation (low glucose, 2.8mM) or 3 hours acute glucose stimulation (high glucose, 28mM) in two independent experiments in INS-1E cells. Distributions are normalized so that the area under the curve equals 1.

B Poly(A) tail length distributions for Chga/b and Iapp upon 1 hour glucose starvation (low glucose, 2.8mM) or 3 hours acute glucose stimulation (high glucose, 28mM) in two independent experiments in INS-1E cells. Distributions are normalized so that the area under the curve equals 1.

C. Scatter plot comparing glucose-induced expression changes (log2 fold change high vs low glucose) in ctrl siRNA versus *Tent5a/c* siRNA treated cells INS-1E cells (n=3, independent experiments). Genes significantly differentially expressed after high glucose (28mM) treatment in control cells (adjusted p-value $p < 0.01$) are highlighted in blue; all others are shown in grey. Differential expression was assessed using DESeq2, after excluding genes with mean normalized counts < 50.

D. Differential gene expression in INS-1E cells following double *Tent5a/c* knockdown and 1-hour glucose starvation (low glucose, 2.8mM), (n=3, independent experiments). MA plot displaying differential expression as log2 fold change (*Tent5a/c* siRNA vs ctrl siRNA) versus mean DESeq2-normalized expression. Genes with adjusted p-value < 0.01 are highlighted in red, while non-significant genes are shown in grey. Selected significantly differentially expressed genes of interest are highlighted in yellow. Differential expression analysis was performed using DESeq2; genes with mean normalized counts < 50 were excluded prior to testing.

E. Differential gene expression in INS-1E cells following double *Tent5a/c* knockdown and 3-hour acute glucose stimulation (high glucose, 28mM), (n=3, independent experiments). MA plot displaying differential expression as log2 fold change (*Tent5a/c* siRNA vs ctrl siRNA) versus mean DESeq2-normalized expression. Genes with adjusted p-value < 0.01 are highlighted in red, while non-significant genes are shown in grey. Selected significantly differentially expressed genes of interest are highlighted in yellow. Differential expression analysis was performed using DESeq2; genes with mean normalized counts < 50 were excluded prior to testing.

**Supplementary table 1.** List of oligonucleotide sequences used in this study.

| Primer | Sequence 5'-3' |
|---|---|
| *Tent5c* fw rat | GATTCTTCGCAGGGTCGTGA |
| *Tent5c* rev rat | GATGAGGTTCAGGGTCTGCC |
| *Tent5a* fw rat | TCGTGCTGGATTGCCTGTTA |
| *Tent5a* rev rat | AGCCCATCAGGTGGCTTTAC |
| *Ins1* fw rat | TACTGCAACTGAGTCCACCA |
| *Ins1* rev rat | TGGTGCTCATTCAAAGGCTT |
| *Ins2* fw rat | AACTACTGCAACTAGGCCCA |
| *Ins2* rev rat | ATTCATTGCAGAGGGGTGGA |
| *Gapdh* fw rat | ATGACTCTACCCACGGCAAG |
| *Gapdh* rev rat | GGTGATGGGTTTCCCGTTGA |
| *Ins1* SP | ATGGCCCTGTGGATGCGCTTCCTGCCCCTGCTGGCCCTGCTCGTCCTCTGGGAGCC CAAGCCTGCCCAGGCTGCTTCCAA |
| *Ins1* 3'UTR fwd cloning oligo | TCGAGGTCCACCACTCCCCGCCCACCCCTCTGCAATGAATAAAGCCTTTGAATGAGC ACCAAaatgagagagtttttatgaatgGC |
| *Ins1* 3'UTR rev cloning oligo | GGCCGCcattcataaaaactctctcattTTGGTGCTCATTCAAAGGCTTTATTCATTGCAGAGG GGTGGGCGGGGAGTGGTGGACC |
